# Automated behavioral profiling reveals baseline phenotypes in marmosets

**DOI:** 10.1101/2024.08.30.610159

**Authors:** William Menegas, Erin Corbett, Kimberly Beliard, Haoran Xu, Shivangi Parmar, Robert Desimone, Guoping Feng

## Abstract

Recent advances in genetic engineering have accelerated the production of nonhuman primate models for brain disorders. To use these models for preclinical drug testing, behavioral screening methods will be necessary to determine how the model animals for each disorder deviate from controls, and whether treatments can restore typical patterns of behavior. In this study, we collected a multimodal dataset from a large cohort of marmoset monkeys and described typical patterns in their behavior and responses to stimuli. We found that these behavioral measurements varied substantially across days, and that behavioral state usage was consistently correlated to the behavior of cagemates and to the total vocalization rate in the housing room. To elicit acute behavioral responses, we presented animals with a panel of stimuli including novel, appetitive, neutral, aversive, and social stimuli. Some stimuli elicited strong and consistent responses, while others failed to elicit large responses. By splitting each animal’s recordings into two halves, we found that stable individual traits could be used to divide animals into groups with test-retest consistency. We found that these groups tend to be shared within families. Finally, we found that some stimulus responses were correlated with baseline behavior in the absence of stimuli, while others were not. This data will help guide the use of marmosets for the preclinical testing of therapies for brain disorders.

## INTRODUCTION

Expanding the number of species across the phylogenetic tree that can be used to test therapies for brain disorders may increase the probability that these treatments can generalize to humans^1,2^. Common marmosets (*Callithrix jacchus*) have recently been established as a nonhuman primate model species^3–6^ that displays several types of social behaviors that are disrupted in human brain disorders, such as vocal communication^6,7^, face processing^8,9^, and social learning^10–12^. We developed high-throughput methods for studying behavior, vocalizations, and responses to stimuli in marmosets. We applied these methods to a large cohort and studied the patterns in data from control sessions and when animals were presented with stimuli. This data will help determine what symptoms are present in genetically engineered model animals, and whether treatments can revert the symptoms^13^.

The application of deep learning to the quantification of video data has recently expanded the potential scope of behavioral measurements. Specifically, methods for extracting animal posture from video data^14–16^ have been transformative because they allow high-dimensional video data (pixels x time) to be distilled into much lower-dimensional postural time series data (keypoints x time) that retains information about the animals’ overt movements^17^. Furthermore, novel approaches to supervised^18,19^ and unsupervised^20–23^ modeling have demonstrated that interpretable patterns in these large new datasets can be identified at many timescales. These computational approaches to ethology have already been applied to the study of genetically engineered mice to reveal similarities and differences across different mouse models of autism spectrum disorder (ASD)^24^. Deep learning has also been applied to the annotation and study of patterns in audio data^25,26^. This has allowed for high-dimensional spectrogram audio data (pixels x time) to be compressed into low-dimensional time series data (calls x time), which can be modeled to understand the basic patterns underlying these complex social processes^27^.

Because the phenotypes of neuropsychiatric disorders are defined in relation to the behavioral patterns of typically developing controls^28^, we collected a large baseline video and audio dataset from group-housed animals living in captivity, comprised of 5,400 sessions from 120 animals living in 36 distinct family units. We found that an animal’s behavioral state usage was correlated with environmental variables such as the behavior of their cagemates and the total vocalization rate in the room where the recording took place. To elicit acute behavioral responses, we presented animals with a set of stimuli including novel, appetitive, neutral, aversive, and social stimuli. We found that patterns in the acute responses to these stimuli were consistent across animals, meaning that this data could be used to benchmark methods for characterizing the response to each stimulus. Finally, by dividing each animal’s recordings into two halves, we asked whether individual differences in behavior were stable over time, whether they were shared within families, and whether baseline behavior in the home cage was related to the responses to stimuli.

Some stimuli caused large and consistent changes in the animals’ position (location) distribution and in the distribution of behavioral states they used, whereas others caused much smaller changes. We found that combinations of baseline and stimulus-response features were stable across the two halves of each animal’s recordings, that animals could be clustered into groups based on these features with test-retest consistency, and that animals tended to be in the same group as their cagemates. Finally, we found that some stimulus responses were correlated with baseline behavior in the home cage, whereas others (such as the approach to the tablet stimuli) were not.

Taken together, these results describe the typical range of marmoset behavior in the home cage and in response to stimuli. We found that the recordings from a single animal can vary as much as the recordings from the whole colony, and that an animal’s behavior is closely related to the behavior of its cagemates. Despite this variability, combinations of baseline and stimulus-response features were stable within individual animals across months of recordings. This study therefore provides a framework for measuring stable individual behavioral profiles in marmosets. These profiles will be important for comparing groups of animals, such as disease models and controls, and for measuring the effect of a treatment within the same animal over time. In both types of experimental design, accounting for the natural variability across days, cagemates, and individual animals could make it easier to detect and interpret the effects of genetic manipulations and treatments.

## RESULTS

### Extracting behavioral data from video

Marmosets form stable breeding pairs that cooperate to raise infants during their development, which takes over a year. Because marmosets often give birth to 2 sibling animals at a time, they usually live in a family of at least 4 total animals when raised in captivity^29^. Because adult females can give birth to multiple litters per year, older siblings need to be paired with new partners in different cages to form their own family units when they are about 15 months old to prevent overcrowding. To study marmosets living in this controlled laboratory environment, we trained a neural network to label 21 postural keypoints of multiple animals (Fig. 1a-b) and developed a classifier to distinguish animals based on the color of the hair tufts near their ears, which we artificially dyed to make them easily distinguishable to a human observer (Fig. 1c-d). This allowed us to extract postural time series data from many different cages with similar family structure (two adult parents and their juvenile offspring).

**Figure 1:**
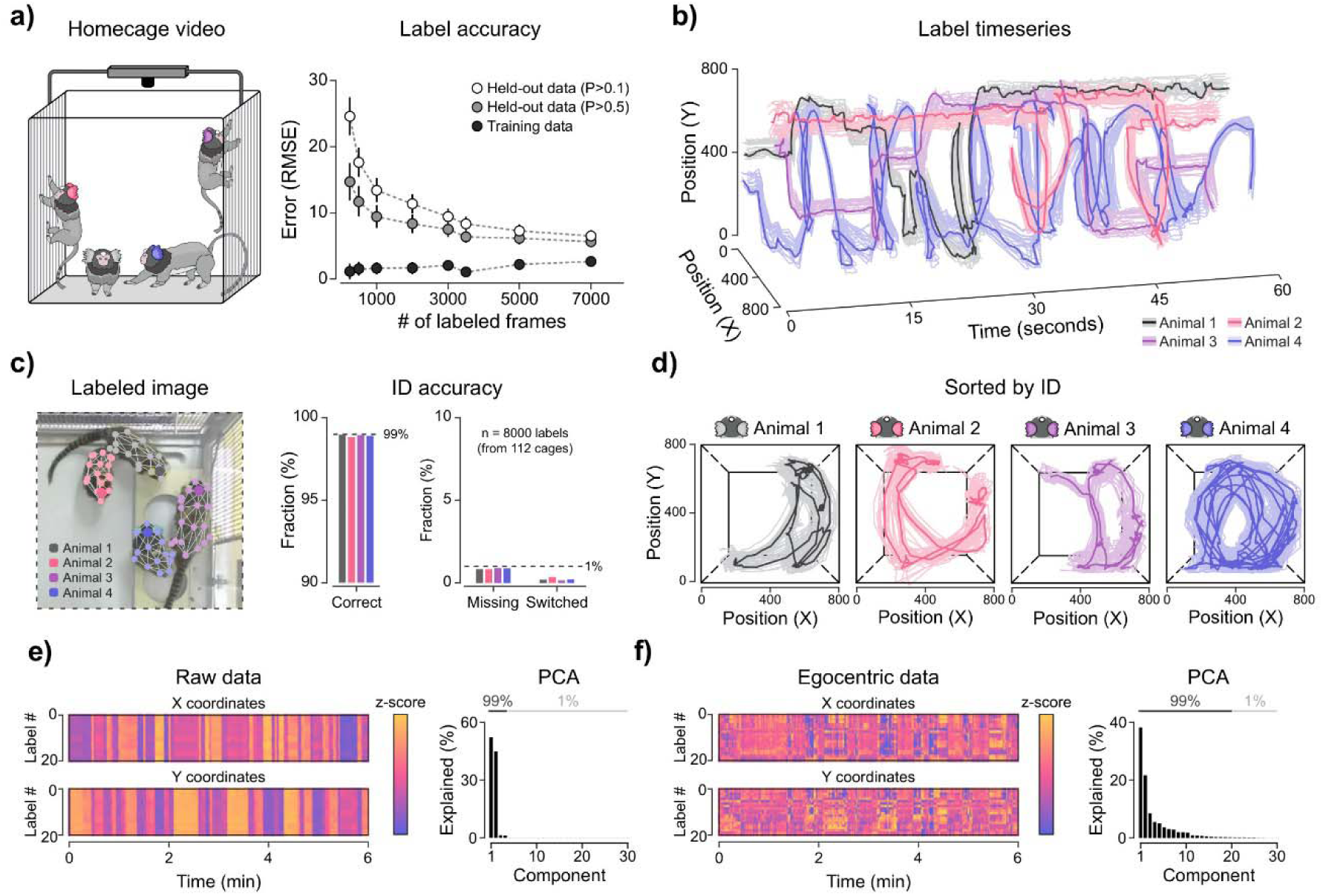
Pose estimation for marmosets in group housing. **a)** Left: schematic of home-cage video recording. Right: keypoint label accuracy reported as the pixel-based root mean square error (RMSE) across an increasing number of labeled frames, for both training data and held-out data (separate lines drawn for two confidence thresholds: P > 0.1 and P > 0.5). Bars indicate the standard deviation across training sessions. **b)** Example time series (60 seconds) of the X and Y positions of all keypoints for the four animals in one recording, colored by animal identity. Thin lines show individual keypoints and thick lines show the head position. **c)** Example labeled image with the tracked keypoints and skeleton of each animal, and ID accuracy based on 8000 labeled animals (one animal drawn per frame) drawn from 112 different camera installations. “Missing” indicates that an animal was not labeled and “Switched” indicates that an animal was labeled with the wrong identity. **d)** Paths of each animal’s keypoints (thin lines) and head positions (thick lines) across 60 seconds, sorted by animal identity. Dashed lines show a schematic of the cage boundaries. **e)** Raw data time series (21 keypoints) across 6 minutes from an example animal (left), and the variance explained by each principal component of the raw data (right). The line above the PCA bars indicates the components that together explain 99% of the variance (4 components). **f)** Egocentric data time series (z-scored) after normalization using X/Y position, orientation, and body size (left), and PCA of the egocentric data (right). The line above the PCA bars indicates the components that together explain 99% of the variance (20 components).

Principal component analysis of the postural time series data revealed that most of the variance was explained by the animal’s X position, Y position, and body angle (Fig. 1e, Fig. S1). This is an expected result from video data in which the animals are small compared to the field of view. To extract more detailed information about the animals’ body movements, we created an egocentric version of the data that was normalized with respect to X position, Y position, body angle, and body size. In this egocentric format, the principal components of variance reflect differences in posture, such as head angle and limb position (Fig. 1f, Fig. S1). Because the Z position of the animal can vary dramatically as animals climb on the cage walls, we used stereo vision to estimate the animals’ depths in the cage (Fig. S2). Recent work has shown that it is also possible to estimate 3D postures directly from 2D postures^30^, and this will be a promising future direction for enabling similar types of high-throughput video data collection with fewer cameras. Additionally, we temporarily installed smooth transparent ceilings (with no handholds) prior to the video recordings to restrict the animals’ ability to climb on the ceiling. We also temporarily placed horizontal cage dividers into the cages to reduce the total cage size during data collection.

Labeling behaviors on a frame-by-frame basis is a difficult task because behaviors can have different timescales and levels of description. Additionally, an animal may simultaneously perform multiple behaviors. Our approach to this problem was to train a recurrent neural network to use long input sequences of features (20 seconds) to predict the following 5 seconds (Fig. 2c). Cagemate behavior was appended to the input to provide context, but the model was only trained to predict the subject animal’s future steps. For paired animals (with only one cagemate), the partner animal was used for context. In families, the same-aged animal was used for context (i.e. the other adult animal or the other juvenile animal). Because the model was trained to predict the subject animal’s future steps, the latent embedding primarily encoded the features of the subject animal (Fig. S4a).

**Figure 2:**
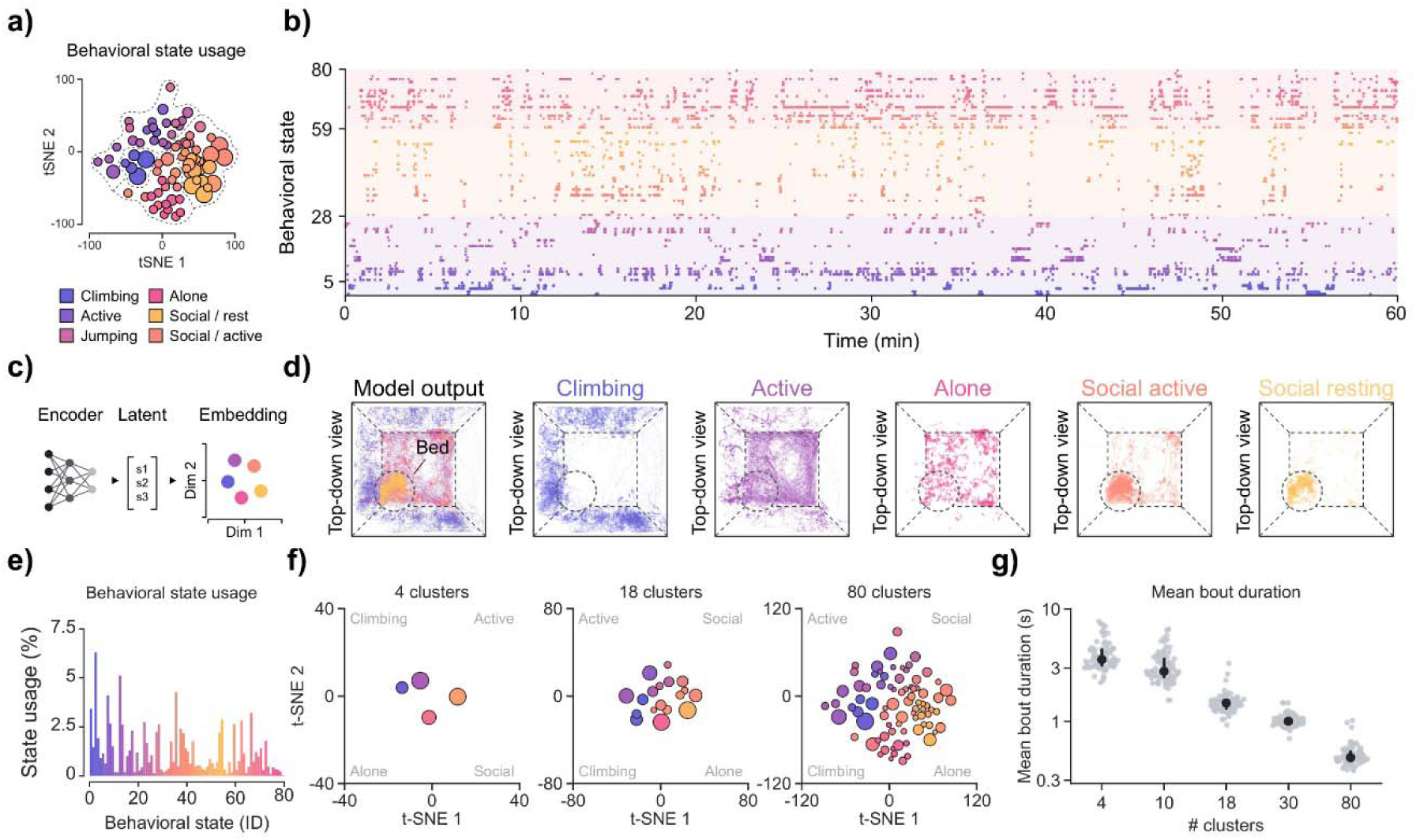
Hierarchical description of marmoset behavior using deep learning. **a)** Average usage of the 80 behavioral states, shown as circles at each state’s position in embedding space, with circle area indicating usage and colors indicating behavioral category. **b)** Example ethogram across 60 minutes from one animal, with each point indicating the behavioral state and colors indicating behavioral category. **c)** Schematic of network structure. The latent state at each time point is used for embedding and downstream analysis. **d)** Top-down view of the spatial distribution of all behavioral categories overlaid (left), and of each of five behavioral categories, pooled over the same 250 sessions (right). The dashed circle marks the position of the bed, and dashed lines show a schematic of the cage. **e)** Average usage of each behavioral state (% of total time) pooled over a random sample of 400 sessions and colored by behavioral category. **f)** Latent space clustering using t-SNE with decreasing values of alpha and perplexity to produce a hierarchical clustering (4, 18, and 80 clusters shown). Circles are plotted at each cluster’s median position, with the circle area indicating usage (% of time) and circle color indicating the cluster identity. **g)** Mean bout duration at each hierarchical level (4, 10, 18, 30, and 80 clusters), with each point representing one animal (n = 86 animals; mean over each animal’s recordings). The median, quartiles, and individual data are plotted.

After training the network, we hierarchically clustered the latent space using t-SNE while systematically varying the alpha and perplexity parameters^31^. We then traced cluster identities across hierarchical levels (Fig. 2f, S3). We found that the finest level of clustering (level 5) resulted in sub-second labels (Fig. 2g) with very specific descriptions (Fig. S4). The coarse level of clustering (level 1) divided the data based on broad categories that humans described as “climbing”, “active”, “social”, and “alone” (Fig. 2a, 2f). The usage rate of these behavioral categories was nonrandomly distributed across physical space in the cage. For example, climbing states occurred most frequently on the cage walls (Fig. 2d), and social resting states occurred most frequently on a bed placed in one corner of the cage (Fig. 2d). The total usage rate of individual states varied widely (Fig. 2e). To increase the human-interpretability of these clusters, we plotted the distribution of features in the embedding space at every hierarchical level (Fig. S4a). This unsupervised hierarchical approach to labeling behavioral states allowed us to study behavioral responses at multiple levels of description.

To understand the temporal structure of the behavioral state sequences, we compared the transitions between consecutive behavioral bouts to sequences in which the order of the bouts was shuffled (Fig. S5a). We found that transitions were concentrated within the same broad behavioral category, and that the observed sequences had a much lower entropy rate than the shuffled sequences (Fig. S5a-b). The mutual information between bouts also remained above that of the shuffled sequences even for bouts more than 200 bouts apart (Fig. S5b). This demonstrates that the behavioral states identified by our model are organized into structured sequences that carry information across timescales, and do not occur in a random order.

### Large-scale data collection reveals variability across days and correlation between cagemates

To treat the study of marmoset behavior as a quantitative data science problem^32,33^, we collected a large dataset of behavioral data from animals living in a laboratory setting. We recorded video data from 120 individual animals living in 36 families at time points when the juveniles were 3, 6, 9, 12, and 15 months old. Some parents were observed with different sets of siblings, as the data collection was spread across several years.

To measure how much behavior varied from day to day, we first examined individual sessions (defined as the data from one animal on one day). Within a single session the relative usage of behavioral categories could switch over the course of minutes (Fig. 3a). The total behavioral state usage also differed significantly across days (Fig. 3b). In total, we recorded 5400 sessions. To compare them, we sorted the sessions along the first principal component (PC1) of behavioral state usage across all sessions (Fig. S7). The sessions of single animals varied widely along this axis (Fig. 3c, Fig. S6). We then compared this animal’s sessions with the sessions of all animals in the colony (Fig. 3d, Fig. S7). The recordings from individual animals spanned nearly the full range of the colony. This observation suggests that the day-to-day variability within one animal could be as large as the variability across all animals. Therefore, a single session of data reveals little about an animal’s behavioral profile, and measuring the stable traits of individual animals could require weeks or months of data per animal. To determine whether stable traits could be measured more efficiently, we next examined the sources of this variability.

**Figure 3:**
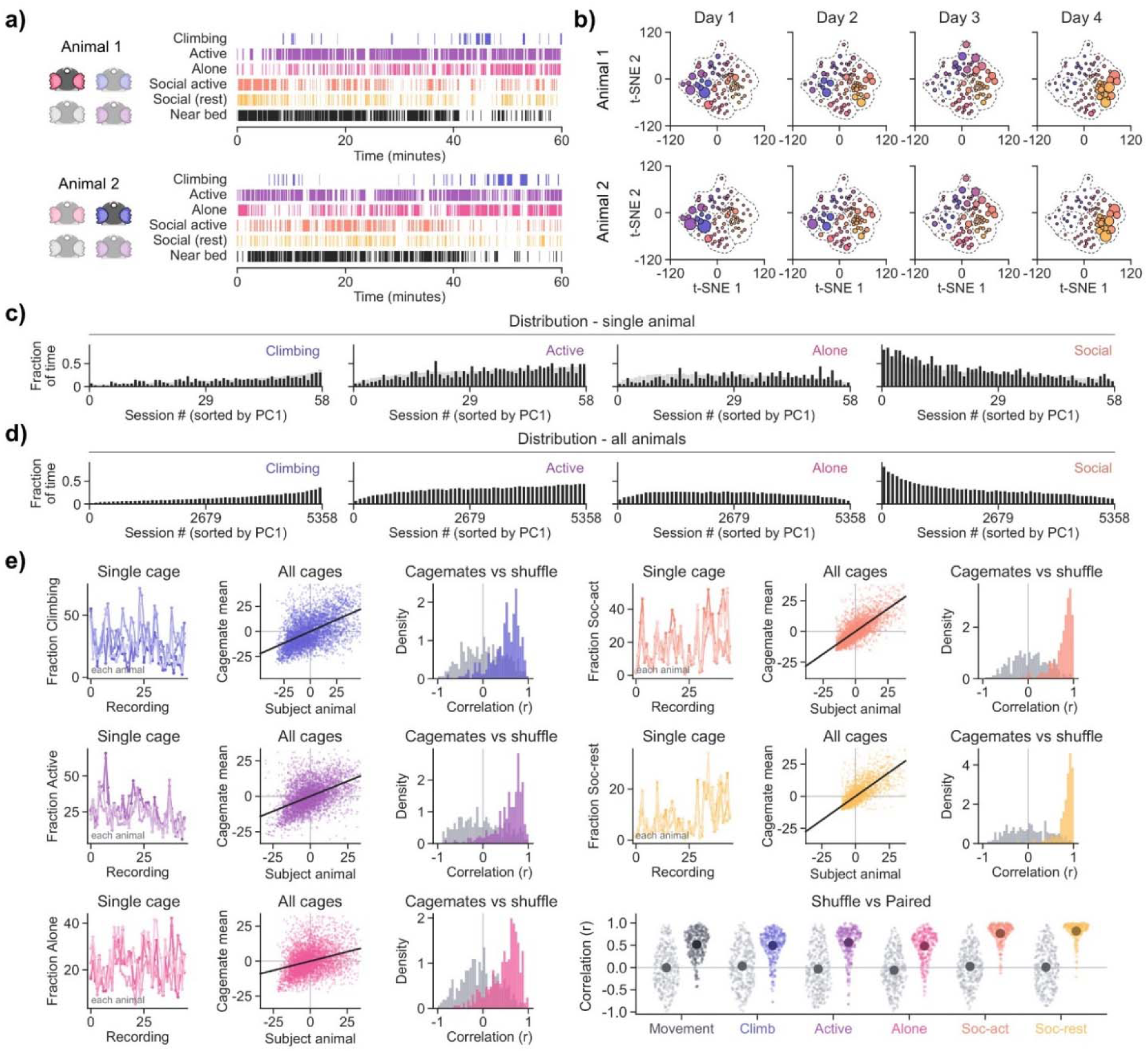
Behavioral distribution varies across days and is correlated to cagemates. **a)** Example time series of behavioral categories and time spent near the bed for two cagemates across one 60-minute session. **b)** Example behavioral state usage from 4 days (non-consecutive) sampled from the same two cagemates. Circle area indicates the frequency of each behavioral state. **c)** Example distribution of behavioral categories across 58 sessions sampled from a single animal, sorted based on PC1 from principal component analysis across all sessions. **d)** Distribution of behavioral categories across 5358 sessions sampled from 116 animals, sorted based on PC1 and averaged in bins of 100 sessions. **e)** Correlation between cagemates within each behavioral category. Single cage: percentage of frames in each category across the recordings of one example cage (one line per animal). All cages: correlation between each animal’s usage of the behavioral state and the mean of the cagemates’ usage of that state. Cagemates vs shuffle: distribution of the correlation between each animal and its cagemates’ mean (colored). For comparison, correlation between the animal and an animal from a different cage and recording period is shown in grey. Shuffle vs Paired: the same correlations for movement and for each category. Correlations with cagemates shown in color and shuffled correlations with an animal from a different cage shown in grey. Darker circles and bars indicate the mean and 95% confidence interval.

When measuring the time spent in each behavioral category across 45 sessions (from non-consecutive days) in an example family, we found that cagemates displayed highly similar patterns of activity^34^ (Fig. 3e). We pooled data from all simultaneously measured animals and found a positive correlation between each animal’s time in each behavioral category and the average time its cagemates spent in that behavioral category in the same session (Fig. 3e). As a control, we compared these correlations to shuffled data. We found that the correlation to cagemates was much higher than the correlation to these shuffled data for every behavioral category, as well as for average movement speed (Fig. 3e). Because social states require two animals to be near each other, some of this correlation is expected for the social categories (since there were only four animals per cage). Overall, these results suggest that measurements of animals’ behavioral traits are strongly correlated to their social environment. Because cagemates are recorded in the same room and with the same recording schedule, external factors such as the call rate in the room could also contribute to this correlation.

### Correlation between total call rate in a housing room and behavioral state usage

Families of marmosets are housed in separate cages, but animals can make loud calls that could influence the behavioral state of neighboring cages despite the lack of visual access^35^. Because we found that animals’ behavioral state usage was correlated to their cagemates’ behaviors, we also wanted to measure the correlation between the rate of these loud vocalizations (that could be heard throughout the housing room) and behavioral state usage. To do this, we installed microphones outside of each cage in a room, such that the signal would be loudest when a call originated from a nearby cage (Fig. 4a, S8). In this setup, all loud calls in the room could be detected by each microphone, and the source could be roughly localized to a single cage. Visual barriers between cages prevented animals from directly observing their neighbors, as is the standard in this colony. We trained a supervised neural network to identify segments of audio data that contained vocalizations using human-labeled training data (Fig. 4b) and trained a separate network to classify the vocalizations according to their call type (Fig. S9).

**Figure 4:**
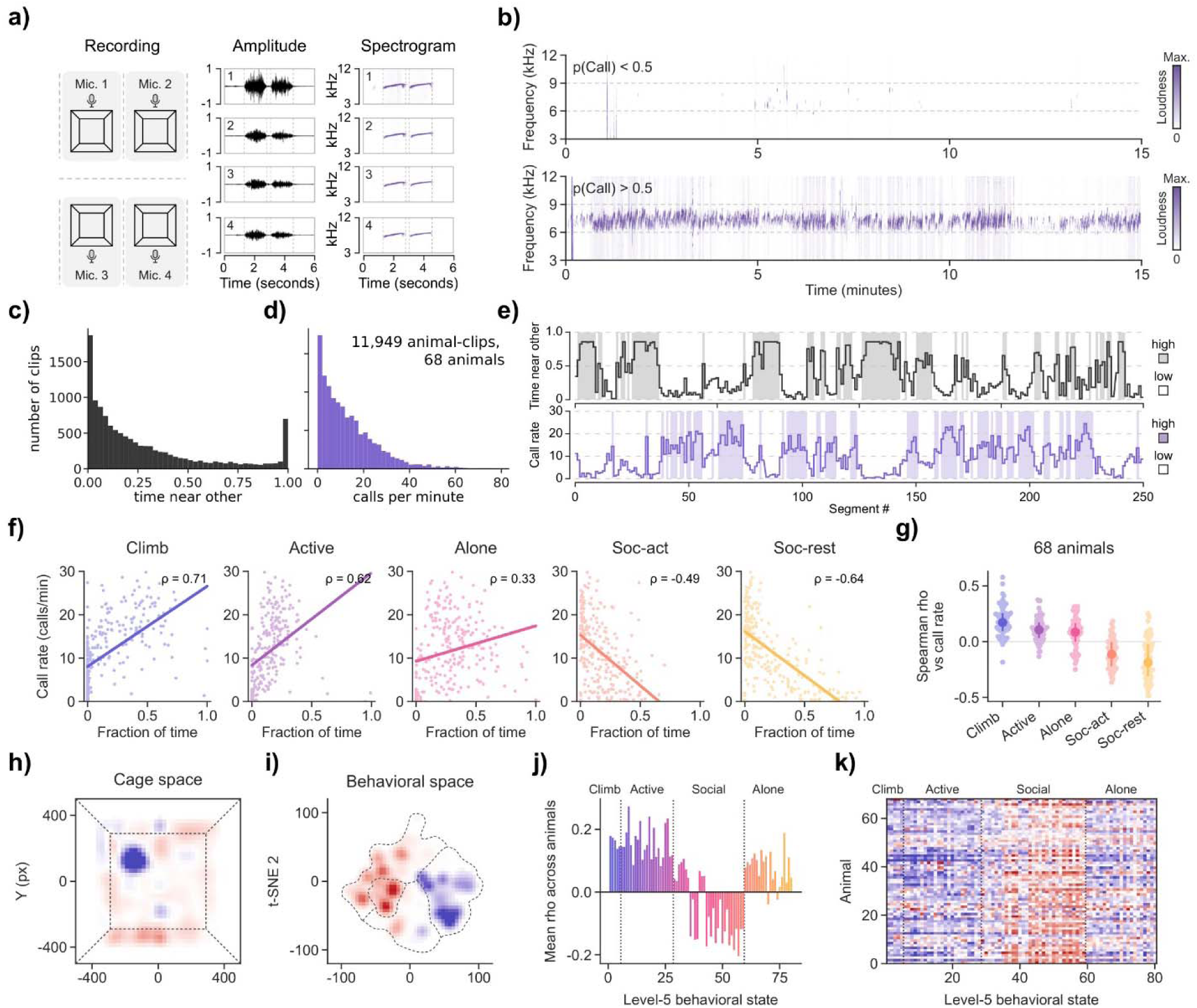
Call rate varies across days and is correlated to behavior. **a)** Schematic of microphone distribution and example raw amplitude data converted to spectrogram data. **b)** Example audio data classified by a supervised neural network as containing no calls (top) or calls of any type (bottom). **c)** Distribution of the time spent in social behavioral states (states 29-59) in 5-minute clips from paired animals. **d)** Distribution of call rates in the same 5-minute clips, recorded by the microphone beside the pair’s cage (n = 11,949 clips from 68 animals). Both of the paired animals’ data was included, for each clip. **e)** Example series of 5-minute clips from one animal comparing the time in social states (top, black line) and the call rate (bottom, purple line). **f)** Correlation between call rate and the fraction of time spent in each behavioral category in an example animal. Each point represents one 5-minute clip (n = 253 clips). ρ indicates Spearman correlation and lines indicate least-squares fits. Data above 30 calls/min are clipped. **g)** Correlation between call rate and behavioral category usage for each animal (n = 68 animals), with each point representing the average Spearman correlation across the states of that category. **h)** Average difference in cage occupancy between 5-minute clips with call rates above and below each recording’s median (n = 56 animals). Red indicates locations that are more occupied when the call rate was high and blue indicates locations that are more occupied when the call rate was low. **i)** The same comparison in behavioral space (n = 50 animals). Dashed lines outline the behavioral categories. **j)** Average Spearman correlation between call rate and the usage of each behavioral state across animals. Dotted lines indicate boundaries between behavioral categories. **k)** Spearman correlation between call rate and the usage of each behavioral state for each animal, with blue indicating positive correlation between behavioral state usage and call rate and red indicating negative correlation.

We first analyzed data from paired animals (see Methods). We split the audio data into 5-minute segments and found that, similar to the time spent in social behavioral states (Fig. 4c), the call rate varied dramatically across segments: from 0 calls per minute to almost 80 calls per minute (Fig. 4d-e). We measured the correlation between call rate and behavioral state usage and found that call rate was negatively correlated with the social behavioral categories and positively correlated with the climbing, active, and alone categories (Fig. 4f-g). When call rates were high, animals shifted their occupancy of the cage (Fig. 4h) and the behavioral embedding space (Fig. 4i), spending less time in social resting states and more time in climbing states. To better understand this effect, we plotted the correlation across all 80 low-level behavioral states (Fig. 4j-k) and found that call rate was most strongly anti-correlated with social behavioral states in which the animals are at rest. This suggests that a high rate of calls in the room could inhibit social resting and cause animals to preferentially exhibit more active/attentive states. Notably, the animals being measured could also be the source of the calls, themselves. So, this effect could also be related to general arousal (in animals that are both making calls and exhibiting more large movements). To confirm the generality of this effect, we repeated the analysis in family-housed animals (typically four animals per cage). In both datasets, a similar overall pattern emerged (Fig. S10), indicating that the pattern was not specific to a particular family organization.

### Measuring behavioral responses to a panel of stimuli

Because the animals’ behavior was highly variable and correlated to cagemate behavior and facility conditions, we aimed to measure species-typical responses that could be robustly elicited by a set of stimuli. We selected a range of stimuli including positive, neutral, negative, novel, and social stimuli (Fig. S11a). These stimuli were presented to the animals over the course of a 3-week recording period, which was repeated every 3 months based on the age of the juveniles in the cage. To better understand this data, we looked for correlations between behavioral state usage and the amount of time spent interacting with each stimulus (Fig. S11b).

As an example of this type of analysis, we compared control sessions to sessions in which a tablet which dispenses liquid reward was presented to animals (Fig. 5a-f). We found that the total usage of most behavioral states was similar between these conditions. Although the conditions were separable based on behavioral state usage, the tablet sessions overlapped substantially with control sessions (Fig. 5d). By contrast, the correlation patterns between behavioral state usage and the time spent near the front of the cage (where the tablet was placed) were very different between conditions (Fig. 5e-f). For example, social behavioral states were positively correlated with the time spent near the tablet in tablet sessions, but not with the time spent at that same location in control sessions. This resulted in a much higher discriminability between the conditions (Fig. 5f). Overall, these results suggest that a stimulus can robustly change the relationship between behavioral state usage and the spatial occupancy map, even when it causes only small changes in the overall usage of each behavioral state.

**Figure 5:**
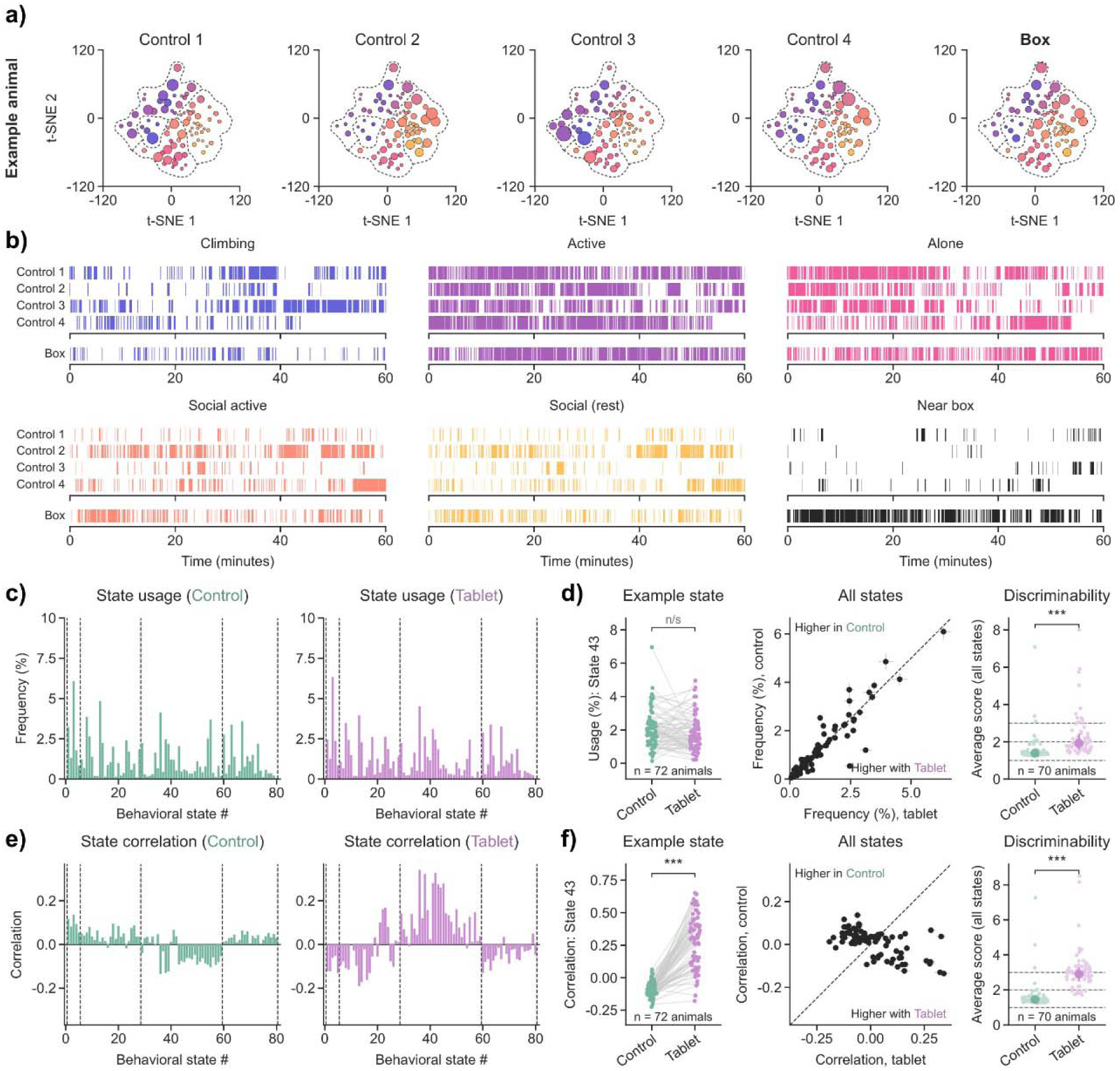
Comparing across behavioral conditions induced by stimuli. **a)** Example behavioral state usage from one animal in four control sessions and one session with a tablet-based stimulus. **b)** The same sessions shown as a time series of each behavioral category and of time spent near the tablet location (black). **c)** Average usage of behavioral states in control sessions (green) and when the tablet is presented (purple). Data was averaged within animals prior to being averaged across animals (n = 72 animals from 229 recording periods). **d)** Example state: usage of state 43 in each animal (n = 72 animals). All states: average usage of each behavioral state in control and tablet sessions. Discriminability: average across states of the absolute deviation of a held-out control session or tablet session from the animal’s remaining control sessions (n = 70 animals). **e)** Average correlation between behavioral state usage and time spent near the tablet location, reported as the Pearson correlation across 1-minute bins of each session in control sessions (green) and tablet sessions (purple). **f)** Difference in correlations between conditions and discriminability between conditions based on these correlations (n = 70 animals). Paired t-tests. * p < 0.05, ** p < 0.01, *** p < 0.001.

### Measuring the strength and consistency of stimulus responses

To compare the responses to all eight stimuli, we computed maps of where each animal spent time in each behavioral category during the first 30 minutes of each session. Most stimuli changed both where the animals spent time in the cage and which behavioral states they used when compared to control recordings (Fig. 6a, Fig. S11c). For example, animals gathered near the tablet (an appetitive stimulus) and moved away from the mask (an aversive stimulus). To measure the size of each animal’s response, we projected the difference between its stimulus and control maps onto the average response of all other animals (Fig. S12). This measurement is positive when an animal responds in the same way as the group. Almost every animal had a positive projection when responding to the tablet, the intruder, the mask, and the treat (Fig. 6b), demonstrating that these responses were highly consistent across animals. By contrast, responses to the toys, noise, and the novel object were smaller and less consistent (Fig. 6b).

**Figure 6:**
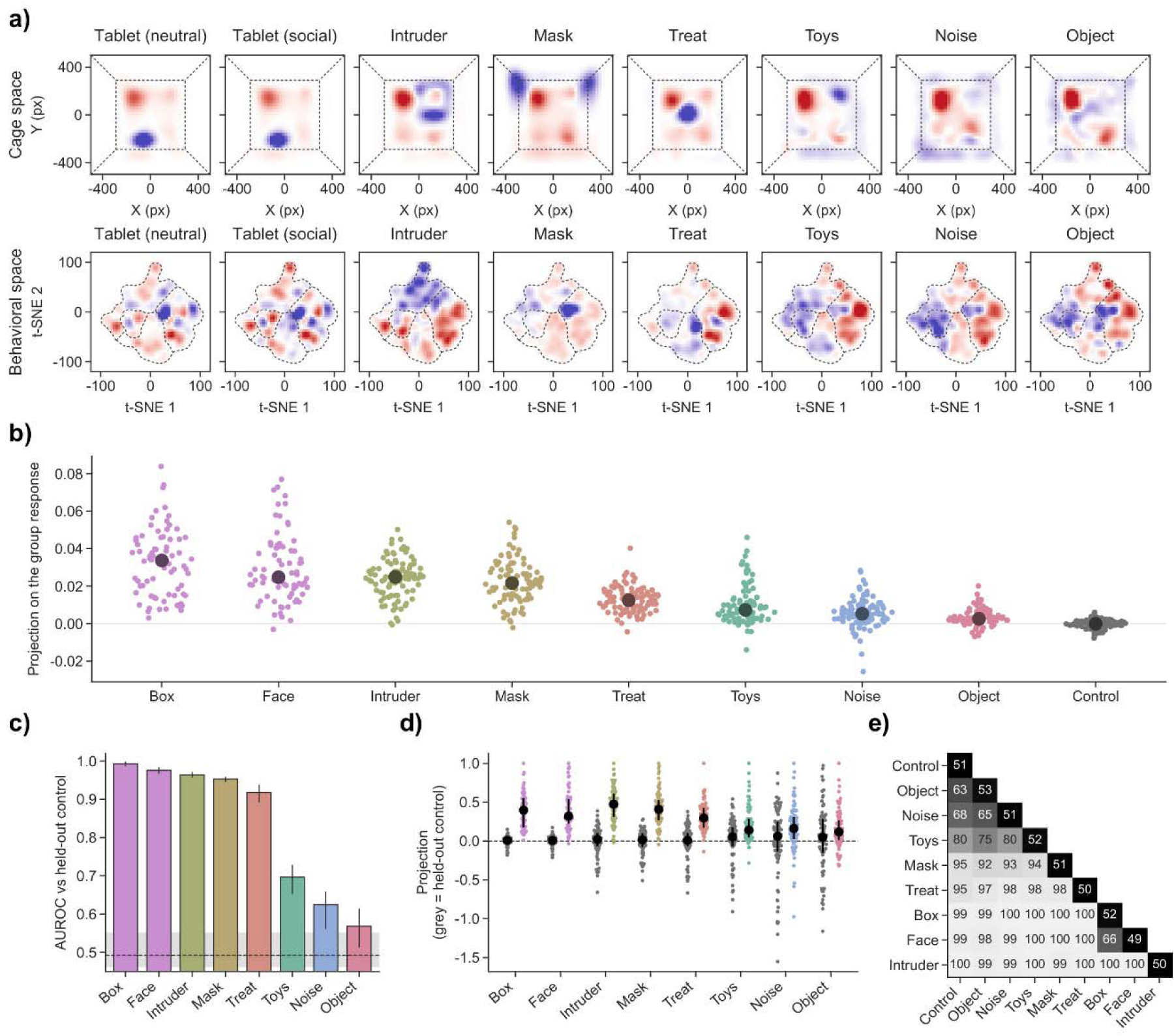
Stimuli induce consistent changes in behavioral patterns. **a)** Difference in physical cage space occupancy (top) and behavioral embedding space occupancy (bottom) between each stimulus and control sessions (over the first 30 minutes). Blue indicates more occupancy in the stimulus session and red indicates more occupancy in the control sessions. **b)** Projection on the group response for each stimulus and for a control session treated as a stimulus (grey), with each point representing one animal and filled circles indicating the median across animals (neutral tablet: n = 72, social tablet: n = 72, intruder: n = 91, mask: n = 92, treat: n = 91, toys: n = 82, noise: n = 92, object: n = 89, and control: n = 92 animals). See Fig. S12 for details. **c)** Discriminability of each stimulus response from a held-out control session (AUROC). Bars indicate the mean and error bars show the range across 10 random draws of the held-out control. **d)** Projection on the group response for each animal in the held-out control session (grey) and the stimulus session (colored), normalized to the largest stimulus projection for each stimulus. Filled circles and bars indicate the median and quartiles (25%, 50%, 75%). **e)** Discriminability (AUROC) between every pair of session types, based on each animal’s projection on the group response. Values on the diagonal compare sessions of the same type.

We could also find similar effects using simpler measurements. For example, animals spent more time near the treat and tablet and spent less time near the mask (Fig. S13a). Furthermore, most stimuli reduced the time spent in social resting states (Fig. S13b). This could reflect an increased state of arousal or interest caused by the stimulus, regardless of valence. Interestingly, the first control session of each recording period also differed from the later control sessions (Fig. S13a-b), suggesting that there is a habituation period where animals adjust to the recording setup during the first session.

We next measured how well each stimulus session could be distinguished from a control session that was held out from the baseline, using the area under the ROC curve (Fig. 6c-d). The tablet, intruder, mask, and treat sessions could be reliably distinguished from held-out control sessions (AUROC > 0.90). By contrast, the toys, noise, and novel object sessions were much harder to distinguish (Fig. 6c). As a negative control, we treated one control session as a stimulus session, which resulted in chance-level discriminability (AUROC ∼ 0.50; Fig. 6c). Comparing all pairs of conditions, we found that most stimuli could be distinguished from each other (Fig. 6e) as well as from control (Fig. 6e). One exception was that the two tablet conditions, which differed only in the image shown on the screen (neutral or social), could not be differentiated from each other.

Also, the weaker stimuli (toys, noise, and novel object) were difficult to distinguish from each other or from control sessions (Fig. 6e).

### Stable behavioral traits and trait groups

Because behavioral state usage varied dramatically across days and was correlated with the behavior of cagemates (Fig. 3), we next asked whether individual animals had stable behavioral traits that could be measured across recording periods^36^. To do this, we split each animal’s recording periods (which were collected every 3 months) into two halves, either by alternating between periods or by comparing the earlier periods to the later periods (Fig. 7a). We then measured a set of 10 features that described the animals’ behavior during control sessions and their responses to the tablets and the mask (Fig. 7d-f). We compared each animal’s features in the first half of its recordings to its features in the second half. For this analysis, we used 64 animals (35 adults and 29 juveniles from 19 families) that had complete data including responses to stimuli from at least two recording periods (see Methods).

**Figure 7:**
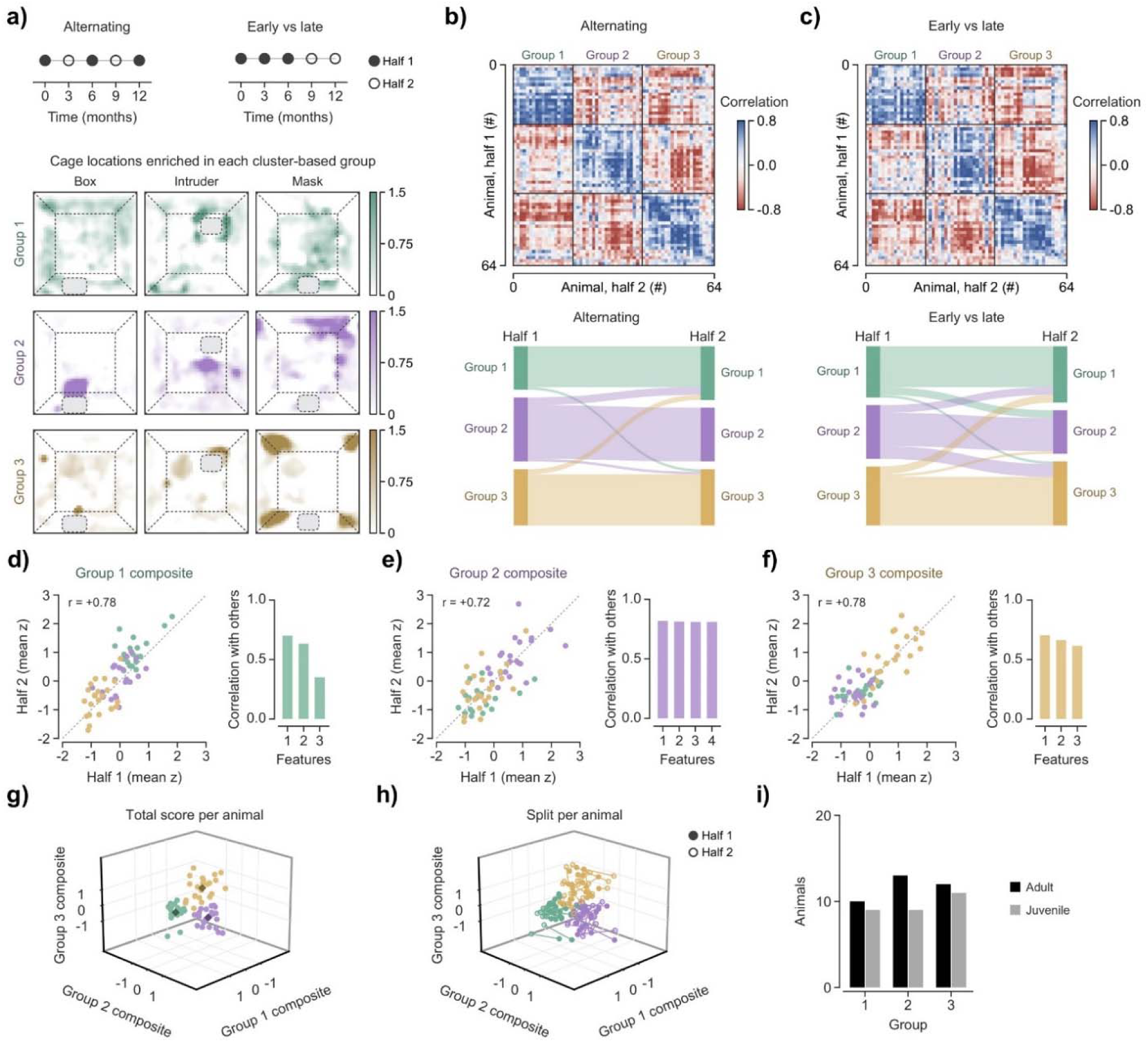
Animals can be clustered into groups based on stable traits. **a)** Data from each animal was split into two halves. Across five recording periods sampled 3 months apart, data was split either into alternating periods (left) or into early/late halves in time (right). Below: cage locations enriched in each group under the neutral tablet, novel animal, and mask stimuli. Color shows the group’s mean response relative to the mean across all animals. White indicates average or below-average occupancy. Dashed lines show a schematic of the cage. **b)** Correlation between every animal’s first half (rows) and every animal’s second half (columns) across the 10 features (for alternating halves). Animals are ordered by group, and the diagonal is a comparison of each animal with itself. Below: each animal’s group assignment in the first half (left) and second half (right), with ribbon showing the transitions across halves. **c)** Correlation between every animal’s first half (rows) and every animal’s second half (columns) across the 10 features (for early/late halves). Animals are ordered by group, and the diagonal is a comparison of each animal with itself. Below: each animal’s group assignment in the first half (left) and second half (right), with ribbon showing the transitions across halves. **d)** Group 1 composite score in the first half and second half (alternating halves), with each point representing one animal, colored by group. r indicates Pearson correlation. Right: The features of the composite, with bars indicating each feature’s correlation with the mean of the composite’s other features. **e)** Same plots for Group 2. **f)** Same plots for Group 3. **g)** Visualization of the three composite scores, with each animal’s mean shown as a point colored by group and diamonds indicating the group means. **h)** Visualization of the same space, with lines connecting the first half (filled) and second half (open) of each animal. **i)** Number of adults and juveniles in each group.

We found that each animal’s features in the first half of its recordings were similar to its own features in the second half (Fig. 7b-c, diagonal). We also found that clusters appeared in the data where subsets of the animals shared similar features across recording periods (Fig. 7b-c). We clustered the animals based on the similarity between their first halves and the other animals’ second halves. By doing this, we divided the animals into three groups (Fig. 7b), that were each characterized by a different combination of features (Fig. 7d-f). Animals in group 1 climbed more during the neutral tablet session and spread across more of the cage and used more behavioral states during the mask session. Animals in group 2 spent more time near the tablets and had larger responses to the tablets. Animals in the third group, which we refer to as the anxious group^37^, spent more time climbing in control sessions. During the mask presentation, they became more active, spent more time active near their cagemates, and spread across less of the cage (spending more time in the upper corners) (Fig. 7a).

We found that group membership was highly consistent across the two halves of the recordings. When we assigned each half to the group whose profile it most closely matched in the other half, both halves were assigned to the same group ∼90% of the time when alternating periods were compared (Fig. 7b), and ∼75% of the time when the earlier periods were compared to the later periods (Fig. 7c). The features that define each group could be combined into composite scores that are highly correlated across halves (r ∼0.75; Fig. 7d-f), and the three groups occupied different parts of the space defined by these 3 composite scores (Fig. 7g-h). Although juvenile and adult animals differed in their overall behavioral state usage (Fig. S14), adults and juveniles were similarly distributed across the three groups (Fig. 7i).

One caveat to this analysis is that the 10 features were selected from a larger set of features (see Methods), and that other selections of features or other numbers of groups could also be used to divide the animals. For this reason, we consider these three groups to be one example of the patterns that can emerge from this type of data, rather than the only way or the best way to divide animals. Overall, these results demonstrate that groupings based on combinations of baseline behavior and stimulus responses are stable within individuals across months of recordings.

### Trait groups tend to be shared within families

Because the animals’ behavior was correlated with the behavior of their cagemates (Fig. 3e), we next asked whether animals living in the same family were frequently assigned to the same groups (Fig. 8a). The two adults in a cage were in the same group in 15 of 16 pairs (Fig. 8a), and juvenile siblings of the same age were in the same group in 12 of 14 pairs (Fig. 8a). Parents and their offspring were also more likely to be in the same group than expected by chance, whereas shuffled pairs of adults from different families or shuffled pairs of juveniles from different families were not (Fig. 8b). The similarity between animals living in the same cage suggests that shared environmental variables could contribute to these groupings^38^. However, because cagemates were recorded together in every session, we cannot fully separate the effect of a shared trait from the effect of being recorded together. One piece of evidence suggesting that this effect was environmental and not primarily genetic is that adult animals in the same cage were not siblings (unlike the juvenile animals), but they still had similar behavioral profiles. If these profiles were primarily genetic, the juvenile-juvenile similarity would have been larger than the adult-adult similarity. Future studies could more thoroughly investigate this by measuring the behavior of animals in their initial family cages and then following them longitudinally after they are placed into new stable pairings.

**Figure 8:**
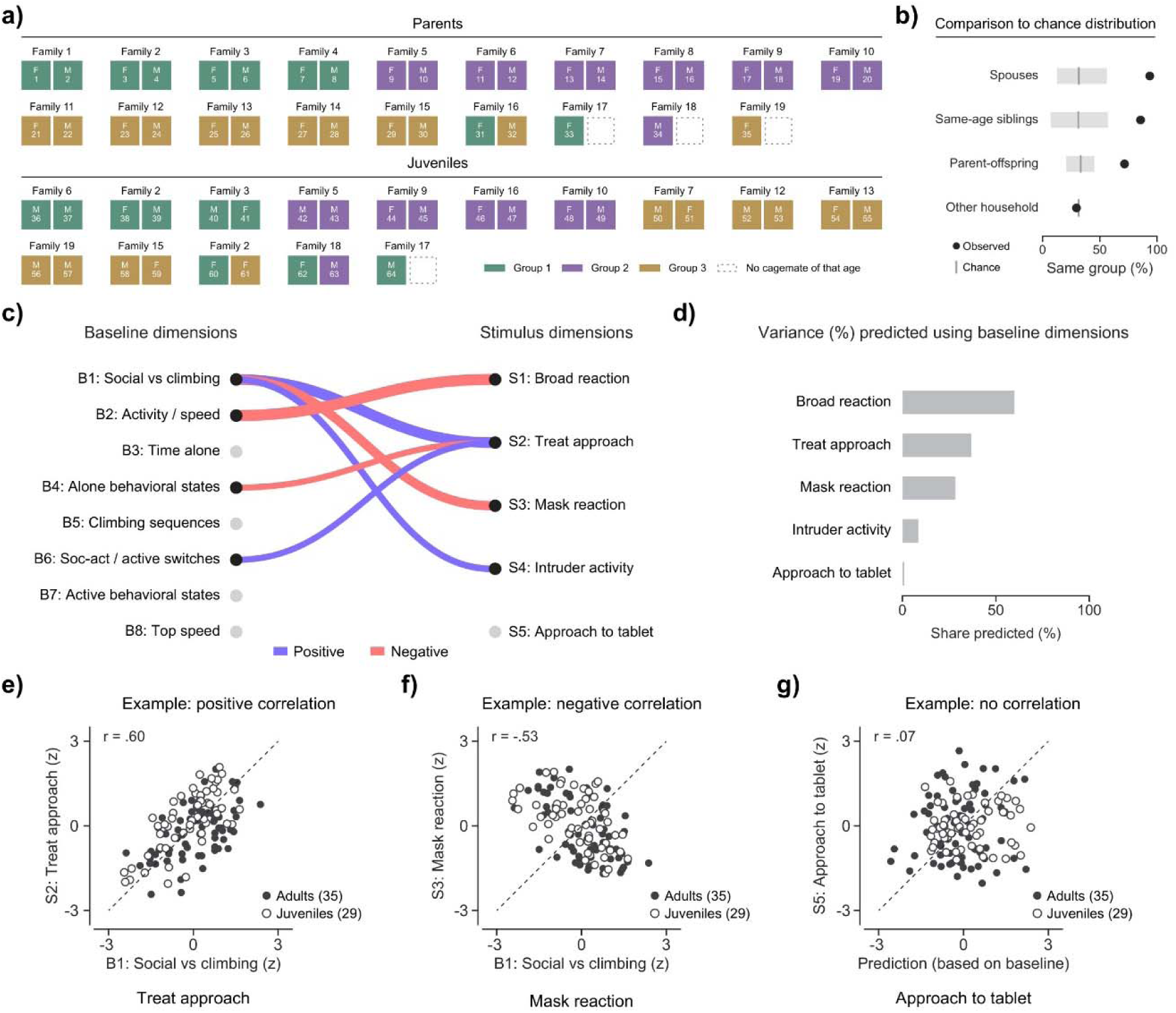
Animals tend to share stable trait groups with their cagemates. **a)** The distribution of groups (as defined in Fig. 7) in each family. Top: adult animals (female left, male right). Bottom: juvenile animals (mixed sex). Each square represents one animal and is colored based on group. Dotted squares indicate spaces where there was no cagemate of that age among the 64 animals analyzed. **b)** Percentage of pairs in the same group for spouses, same-age siblings, parent-offspring pairs, and same-age animals from other families. Grey line indicates chance distribution. **c)** True-score correlations between the 8 baseline dimensions (B1-B8) and the 5 stimulus dimensions (S1-S5). Lines join pairs whose true-score correlation is at least 0.4 in absolute value (blue, positive; red, negative). Line width is proportional to the absolute value of the correlation. See Methods for details. **d)** Fraction of each stimulus dimension’s reliable variance predicted from all baseline features by nested ridge regression in held-out families. **e)** Baseline dimension B1 is positively correlated with the stimulus dimension related to the approach to the treat (Pearson correlation). **f)** Baseline dimension B1 is negatively correlated with the stimulus dimension related to the reaction to a mask (Pearson correlation). **g)** Approach to the tablets is not predicted by baseline features in held-out families. All panels use 64 animals from 19 families, with each animal’s recording periods divided into alternating halves.

### Baseline behavior is correlated with some stimulus responses

Finally, we asked whether the animals’ responses to stimuli were related to their baseline behavior in the home cage. We measured 1,337 baseline features in control sessions and found that 790 of them had test-retest reliability, meaning that they were correlated across the two halves of the recordings. We grouped these reliable features into 8 baseline dimensions. We also grouped the reliable stimulus-response features into 5 stimulus dimensions and found that some baseline dimensions were predictive of the stimulus-response dimensions (Fig. 8c). For example, animals that spent more time in social states and less time climbing at baseline (baseline dimension B1) scored higher on the treat approach dimension and had smaller responses to the mask (Fig. 8c, e-f).

Using all of the baseline features, we could predict a substantial fraction of the reliable variance in the reactions to stimuli, the treat approach, and the mask reaction in families that were not used to train the model (Fig. 8d). By contrast, the approach to the tablets could not be predicted from baseline behavior (Fig. 8d, 8g), and none of its correlations with the baseline dimensions were greater than 0.4 (Fig. 8c). Overall, these results suggest that some stimulus responses are related to the animals’ baseline behavior, while others capture aspects of behavior that are not predicted by baseline behavior, such as the approach to the tablets.

## DISCUSSION

### Methods for quantifying behavior

Finding patterns in behavioral data is a complex problem, and several different approaches may yield meaningful descriptions of video data. Broadly, the analysis of video data can be either pixel-based^20,21^ or keypoint-based^39,40^. We chose to use a keypoint-based method because of the large variety of cage components, objects, stimuli, and other animals present in the study. Pixel-based methods may be more appropriate when the conditions are more constant across video frames, such as in the quantification of facial expression in head-fixed mice^41^. Another key decision is whether to use supervised or unsupervised learning approaches. Supervised learning can be beneficial in cases where a small number of human-recognizable behavioral modules are the subject of the investigation, such as in studies of aggressive responses^42^ or escape responses^43^ in mice. Unsupervised learning can be more difficult to implement because the patterns identified by the model may not have much biological relevance. For instance, models using raw x/y/z coordinates rather than egocentric coordinates may label different positions in the cage as key behavioral states, rather than different motions. In cases where input data contains missing values or large jumps, the model may identify these features as the main sources of variance. We chose to use an unsupervised learning approach because our goal was to describe the entire distribution of the animals’ daily behaviors in a relatively unbiased manner rather than selecting human-identifiable behaviors manually and quantifying the frequency of those behaviors.

Aside from questions regarding methodology, the study of behavior is also complicated by the lack of a specific definition of what constitutes a behavioral module. Importantly, behaviors can occur on different timescales, and it is not always clear (a priori) which timescale will be relevant for each study. For instance, some model animals meant to display brain disorder symptoms may have motor skill deficits requiring a detailed kinematic analysis of sub-second behaviors^44^ and others may have deficits in complex social interactions that occur across several seconds or minutes. Many studies have focused on sub-second units of behavior because they are the shortest segments that can be clearly differentiated into units, often referred to as syllables^20,21^.

Another complication is that behavior can be described at many levels of specificity. At higher levels of specificity, more data is needed to find patterns. But at lower levels of specificity, patterns may not be apparent due to variety within a single category. We chose a method where each frame would have a single label, while preserving hierarchical information. This allowed us to identify patterns at multiple levels of description. Notably, our recordings were made at 60 frames per second, which allowed us to collect and store data from a large number of cages. This may be sufficient for behavioral states lasting hundreds of milliseconds or longer, but higher frame rates would be needed to precisely resolve the kinematics of all marmoset movements.

### Large-scale data collection and reproducibility

Logistical constraints and ethical considerations limit the ability to collect large behavioral datasets from nonhuman primates (NHPs)^45^, unlike in several other model organisms. NHP research is comparatively much more expensive due to the larger living spaces required, longer generation times, and more complex veterinary care needed to maintain high-quality living conditions. We opportunistically collected data from a large colony of marmosets that was primarily used for transgenic model production. This study will be useful to guide future work to ensure logical experimental design principles for preclinical testing in marmosets.

When comparing behavioral data across facilities, challenges can arise due to environmental variables such as the behavioral distribution of family members, number of vocalizations in the room, room configuration, and stimuli present^46^. Although these conditions may change the exact distribution of behavioral state usage, we expect that the general principles for marmoset behavioral analysis identified in this study should apply across facilities. Importantly, we also found that animals in the same family tended to be assigned to the same trait groups, which emphasizes the idea that social environment should also be considered when comparing animals across cages and facilities.

### High-dimensional phenotyping for neuropsychiatric disorders

Neuropsychiatric disorders can have a wide range of symptoms, and disease model animals will need to be tested in a variety of contexts to study their behavior, responses to stimuli, cognitive ability, and performance in other assays. For example, people with autism can have a variety of symptoms, such as differences in social behavior, vocal communication, cognitive ability, responses to novel or high-intensity sensory stimuli, restricted interests, or repetitive behavior^47^. Some of these symptoms may only manifest in certain conditions, such as in novel environments or when meeting new people. Some symptoms may also be secondary effects, such as differences in organ health^48^. To test which phenotypes are present in disease model animals, high-throughput and high-dimensional methods will be essential. This is particularly important for testing whether treatments can rescue multiple deficits or only a subset of the deficits.

### Stable traits for individual phenotyping

Human patients with ASD can display diverse symptoms^49^ including differences in social affect, repetitive behaviors, restricted interests, lower verbal ability, and impaired cognitive ability. Importantly, this diversity is observed even in monogenic forms of ASD, such as in Phelan-McDermid syndrome^50–52^, which can be studied in animal models with alterations in the Shank3 gene^53,54^. These patients have a range of symptoms, and individual patients can exhibit different combinations of symptoms. For this reason, it will be important to phenotype animals on an individual basis by comparing data from each animal to the whole-population distribution^55^ rather than only on a group basis (comparing across pre-defined groups of animals). This requires knowing which behavioral measurements are stable within individuals over time, and how much these measurements vary across typically developing animals^56^.

Many studies of model mice for brain disorders have focused on group-based differences in specific assays^53,57^. Although bulk differences between heterogeneous classes can be detectable with large sample sizes (as is typical in mouse studies), this is logistically impractical for NHP research. Rather than producing large cohorts of control animals for each study, a more efficient use of animals will be to collect large datasets of control data that can be used to power future studies with smaller cohorts. This could be done for data related to behavior, responses to stimuli, cognitive tests, and other assays. These datasets could be standardized and shared across institutes, as is becoming more common for mouse research^58^. The normative models from these datasets could be used to test whether smaller cohorts of animals (controls, disease models, or treated animals) differ from typical animals in the dataset^55^.

We found that animals could be divided into groups based on stable combinations of baseline behavior and stimulus responses, and that these groups tended to be shared within families. We do not consider these groups to be discrete types of animals, since other selections of features and other numbers of groups could also divide the animals. However, the stability of these groupings across months, together with the similarity between cagemates, suggests that individual differences in marmoset behavior reflect both stable traits of each animal and the social environment. Because family members were recorded together, future studies that record animals separately or after changes in housing will be needed to separate these factors.

We also found that some stimulus responses (such as the approach to the treat and the reaction to the mask) could be partially predicted using baseline behavior while others (such as the approach to the tablets) were not. This suggests that stimulus assays can measure traits that are not fully captured by home cage recordings, and that combining baseline recordings with a panel of stimuli will provide a more complete description of each animal than either approach alone.

## METHODS

### Animals

For this study, we opportunistically collected behavioral data from 120 animals that were part of a large colony used for transgenic animal production. No animals were obtained or bred specifically for use in this project. The animals were all housed in family units consisting of 2 adult animals and juveniles up to the age of 15 months. After 15 months, the juveniles would be paired with other animals in new cages. Data could then be collected from those paired animals which would then become the “adult” generation of their new cage. The lights in the facility were turned on at 7:00 am and turned off at 7:00 pm. Veterinary checks and other work in the rooms (including cage changes) were done between 7:00 am and 10:00 am. Each session of data was collected between 10:00 am and 4:00 pm. After 5:00 pm, the animals were not disturbed to ensure that their sleep cycle would not be affected. The facility was maintained at 26 °C and 45 ± 10% relative humidity.

All procedures were performed in accordance with the National Institutes of Health Guide for the Care and Use of Laboratory Animals and approved by the MIT Committee on Animal Care (CAC). The MIT Division of Comparative Medicine (DCM) routinely monitored animal welfare and provided the animals with veterinary care as needed.

### Home cage video recording

Cages were scheduled for recording periods based on the ages of the juvenile animals living in the cage, such that they would be recorded for 3 weeks at a time when the animals were 3 months, 6 months, 9 months, 12 months, and 15 months old. Recording sessions were scheduled for weekdays to avoid variance in environmental conditions between weekdays and weekend days. Once juveniles were 15 months old, they were transferred to new cages to begin their own families with new partners. Our methods could not be used until juvenile animals reached 3 months old, because that was when the hair tufts near their ears were long enough to dye (to differentiate between animals) and also because the animals began to behave more independently from their parents (rather than clinging to one of their parents) at this time. During the week prior to each recording period, cages were changed to accommodate a transparent ceiling panel to facilitate top-down video recording. The smooth transparent ceiling panels prevented animals from climbing on the ceiling during the recording sessions, and horizontal dividers were added to further restrict the total cage size during the recordings. After these cage modifications, the final size of the area used for the recording was approximately 75 cm x 75 cm x 75 cm.

Prior to the first recording session, cameras were placed above the cages, the ceiling was replaced with a transparent panel, and the animals’ hair was dyed. The hair colors were planned such that each cage would have a red-haired animal, white-haired animal, blue-haired animal, and yellow-haired animal. Red, blue, and yellow were chosen as the dye colors to maximize distinguishability between animals, according to human observers. Human distinguishability is essential to this approach since we used a supervised model to classify animals based on hair color. To apply hair dye, one experimenter caught and restrained the animal while a second experimenter applied the dye into the hair. Dye was applied liberally to the tufts of hair near the ears so that there was no white hair remaining. Animals were then placed in a temporary cage for 20 minutes so that the dye could dry before they were transferred back to their home cage. This was done to prevent family members from rubbing the dye onto themselves and to prevent the dye from marking items and surfaces in the home cage. After animals were returned to their cages, we waited at least one day before any recordings were performed to allow the dye to dry completely and allow the other animals to habituate to the dye.

Video recordings were done using ZED stereo cameras mounted above the cages and collected using the ZED SDK developed by the manufacturer. Video was collected from 2 cameras at 60 frames per second. The image resolution was 720p, meaning that each image had a size of 1280 pixels by 720 pixels. Camera settings (i.e. brightness/contrast/exposure/gain) were manually adjusted after each camera installation to avoid image quality issues caused by oversaturation or insufficient gain. Because light sources directly above the cages could cause glare in the video, we either moved the cages or moved the light sources to ensure that cages were not directly below light bulbs. After exporting the proprietary “svo” videos generated by the cameras into the “avi” format using the ZED SDK, further processing was done using Python and MATLAB.

### Pose extraction

Postural keypoints were extracted from the raw video data using DeepLabCut (DLC)^14^ and sorted using a two-step method we developed for this specific use case. First, poses were joined into tracklets based on kinematic features. Then, these tracklets were sorted based on the average distance between each tracklet and the labels tracking each hair color. To train a network to recognize hair color, we placed one keypoint label on the center of the head of each animal, rather than on each hair tuft. Overall, this approach leveraged kinematic information to join nearby points in x/y/time coordinates and used the hair color tracks to ensure that animal identities were not mixed after animals had close interactions or were occluded by other animals. Egocentric postures were calculated by setting the middle point on the body to (0,0) and then rotating all points based on the angle formed by the front point on the body and the back point on the body. Egocentric data was then normalized by body size. Features were calculated based on the raw time series data, depth information, and the egocentric time series data.

### Behavioral state labeling

A recurrent neural network was trained to use 20 seconds of input data (features) to predict the next 5 seconds of data (features). This network had a bottleneck layer of 256 units. The latent space was the 256-dimensional output of this bottleneck layer, computed for each frame. The data from this bottleneck layer was used for hierarchical t-SNE clustering. As suggested in tree-SNE^31^, we systematically varied the alpha (exaggeration) and perplexity (number of neighbors) in the t-SNE clustering algorithm. At high levels of alpha/perplexity, we observed fewer distinct clusters. At lower levels of alpha/perplexity, we observed more distinct clusters. We refer to each of these clustering results as a hierarchical level, from level 1 (4 clusters) to level 5 (80 clusters). Initial cluster assignment was done using Density-Based Spatial Clustering of Applications with Noise (DBSCAN)^59^. Cluster labels were assigned to new data (obtained after initial model training) using openTSNE^60^ to embed new points into the existing embedding. We then used k-nearest neighbors (KNN) to assign a state label. This was done at the lowest hierarchical level only, because higher levels could be inferred from this information. In addition to the four categories of the highest hierarchical level, some analyses used five behavioral categories (climbing, active, alone, social active, and social resting) based on human categorization of the 80 fine behavioral categories. These five categories are not a subdivision of the four highest-level categories.

### Home cage audio recording

Microphones were placed outside of each cage under study and also outside of each neighboring cage. Small wireless microphones were used so that they could be placed around the room without adding any wires or large equipment that could potentially draw the animals’ attention. Audio data was recorded at 48,000 samples per second (48 kHz) and logged locally on each wireless recording device. The data from each device was synchronized to the video data using a series of 5 claps from a clapper board at the beginning and end of each video. These claps were clearly visible in the video and easily identifiable in the audio data by a human observer. The clap onset time was manually annotated for each video and audio recording.

### Vocalization labeling

Audio data was converted into spectrograms using SciPy^59^ with nperseg = 700, noverlap = 525, and nfft = 2000. These spectrograms were segmented using a supervised neural network trained to perform binary classification where 0 = no call and 1 = call. To determine the average call rate (Fig. 4), a threshold of 0.5 was applied to the output of the neural network, such that all outputs below 0.5 were set to 0 and all outputs above 0.5 were set to 1. Spectrograms were segmented into discrete calls using this binary classification. Next, the discrete calls were classified by a separate neural network trained to perform classification based on human categorization of calls in the training set into 8 classes. The types of calls labeled by this model were limited to the types of calls that were commonly observed in our training data.

### Stimulus presentation

Stimuli were presented over the course of 3 weeks, and the same series of stimuli was presented to each cage every 3 months. Each session began at approximately 10:00 am, and its total length varied with the type of stimulus. Because every session started at the same time of day, we did not analyze changes in behavior across the time of day. To compare sessions of different lengths, the analyses of stimulus responses (Fig. 6, Fig. 7, Fig. S11-S13) used the same window at the start of each session (the first 30 minutes). The remainder of each session (which was slightly variable in length, depending on the schedule and number of animals recorded each day) did not affect these comparisons. The first 5 sessions were “control” sessions. In this condition, a horizontal divider was placed in the middle of the cage to restrict the total cage size and simplify analysis. Ledges, branches, and hammocks were removed so that animals would not be occluded from view. Nest boxes and other cage attachments were also removed, so that animals could not leave the field of view. A bed was placed in the rear-left corner of the cage so that animals had a comfortable place to sit when resting. Food was placed into two bowls that fit into slots designed to hold them so that they remained in place. Water was freely available through spouts on the side of the cages. At the beginning of each session, cameras were turned on and audio data was synchronized to the cameras as described above (using a clapper board to synchronize the audio data to the video). After the completion of each session, cages were restored to their normal state.

After the 5 days of these control sessions, stimuli were presented once per day. During these sessions, the cages were set up as described above and then one of the 8 types of stimuli was added. Including the control sessions, this means that there were 9 total stimulus-response conditions (one control condition and 8 stimulus conditions). Because the control condition was performed 5 times per set of recordings, and the toy condition was performed 3 times per set of recordings, there was not an even number of each condition in the total dataset. The conditions are described in the following section.

*1) Control*. The only items present in the cage were a bed that animals could rest in, two bowls of food placed in slots designed to hold the bowls, and water-dispensers. These items were also present in the other stimulus conditions.
*2) Novel object.* A large and brightly colored plastic dog toy was placed in the cage. Importantly, this type of object is not typically placed in the cage. Marmosets cautiously approach these objects but have low overall interest in them.
*3) Treat*. A mixture of foraging materials and foods high in sugar was placed in a bowl. Foods such as banana chips, marshmallows, pudding, dried cranberries, and dried mango were mixed into the foraging materials. Importantly, these foods were not regularly given as part of the standard diet. Marmosets quickly approach and consume these highly appetitive items.
*4) Noise.* A movie was played on a computer outside the cage. Marmosets did not have visual access to the movie and only heard the dialogue and soundtrack. This “background noise” stimulus is not meant to have a large impact on their behavior.
*5) Mask.* A gorilla mask was placed outside the cage in the same position as the computer playing the “background noise” stimulus. The mask stimulus is similar to “human intruder” experiments^37^. Marmosets have a very large negative response to this stimulus, often climbing to the furthest corner of the cage in a group, and sometimes making angry calls.
*6) Toys.* A set of colorful wicker balls that animals frequently chew on and interact with was placed in the cage. The toys were presented for 3 consecutive days. The wicker balls were attached to the side of the cage using zip ties so that they could not be moved to different positions in the cage, which could complicate the interpretation of the data.
*7) Intruder animal (“Intruder”).* A novel animal from a different family was placed in a transfer box inside the cage. Animals were allowed to have auditory/visual access to the animal but were not able to physically interact (they were separated by a transparent plastic panel). This was done to prevent the injury of either the family or the intruder animals. This is a highly salient stimulus and causes a large increase in activity.
*8) Tablet 1 (neutral tablet).* Animals were presented with a neutral stimulus (blue square, or “box”) in the center of the screen. They could touch the screen to receive a syrup reward that was delivered from a metal spout attached to the bottom of the tablet.
*9) Tablet 2 (social tablet).* Animals were presented with a social stimulus (marmoset face, or “face”) in the center of the screen. They could touch the screen to receive a syrup reward that was delivered from a metal spout attached to the bottom of the tablet.

### Transition structure

Behavioral state sequences were converted into sequences of bouts by merging consecutive frames with the same state. The entropy rate of each session’s bout sequence (the conditional entropy of the next bout’s state given the current bout’s state) and the mutual information between bouts separated by increasing numbers of bouts were compared to sequences in which the order of the bouts was shuffled. These shuffles preserved the total usage rate of each state but randomized the order. Values were averaged over each animal’s control sessions. Stimulus sessions were not included in the analysis for simplicity. Shuffled entropy rates were compared using a paired t-test.

### Correlation between cagemates

For each recording, we calculated the fraction of time animals spent in each of the five broad behavioral categories. We also calculated the average movement speed of each animal. To normalize this data, we subtracted the colony mean for that session type from each value. Within each cage and recording period, we then correlated each animal’s values with the mean of its cagemates’ values across their shared recordings. Only animals with at least 5 shared recordings were included. As a reference, we measured the correlation of each animal’s values with the cagemate mean from one randomly chosen recording period of another cage, or of the same cage at another time.

### Call rate and behavior

Audio and behavioral data were divided into 5-minute segments, and the call rate in each segment was calculated from the microphone beside the animals’ cage. For each animal, we calculated the Spearman correlation between the call rate and the usage of each behavioral state across segments and averaged these correlations across the states of each behavioral category. To compare the occupancy of the cage and of the behavioral space between high and low call rates, we split each recording’s segments at that recording’s median call rate and averaged the difference between the high and low maps across animals.

### Comparison of control and tablet sessions

For each animal with at least two control sessions in a recording period, we compared two measurements between control sessions and neutral tablet sessions: (1) the usage of each behavioral state, and (2) the correlation between the usage of each state and the time spent near the tablet location. We calculated this correlation using the Pearson correlation across data chunked into 1-minute time bins. For control sessions, the time spent near the same location in the cage (without the stimulus being present) was used to calculate these correlations. To measure discriminability, we calculated the absolute deviation of a held-out control session or of the tablet session from the animal’s remaining control sessions for each state, in units of standard deviation. We then averaged this metric across states. Control and tablet-response conditions were compared using paired t-tests.

### Stimulus response maps

For each session, we divided the first 30 minutes into 0.5-second time bins. For each time bin, we recorded the animal’s head position (using 20-pixel spatial bins) and its behavioral state. Each behavioral state was assigned to one of five behavioral categories (climbing, active, alone, social active, and social resting). We counted the time bins in each category and location to create five maps (5 categories × 41 × 41 locations). Each map was normalized by the total number of time bins in the session and smoothed with a Gaussian kernel (σ = 1 spatial bin). We included recordings with at least 60,000 frames and sessions with at least 500 bins. For each animal, the baseline control map was defined using the average of that animal’s control sessions from all recording periods. One control session per animal was held out at random and was not included in these baseline maps. We defined the animals’ response as the difference between the stimulus map and the baseline map.

To measure the size of the animals’ responses, we projected each response onto a template (Fig. S12). The template was defined using the average response of all other animals. The animal’s own data was not included in its template, so the template was slightly different for each animal. We also subtracted the baseline map from the held-out control session and projected it onto the same template.

Discriminability was measured as the area under the ROC curve (AUROC), comparing the stimulus projections with the held-out control projections across animals. In this metric, a score of 0.5 would indicate no discriminability, and a score of 1.0 would indicate perfect discrimination. We repeated this procedure with 10 random draws of the held-out control session, and each animal’s projection was averaged over these draws. As a negative control, we treated one control session as a stimulus and found no discriminability. For comparisons between pairs of conditions (Fig. 6e), we projected each animal’s average map in each condition onto the average difference between the two conditions in all other animals. The AUROC then compared the two sets of projections.

For Fig. 6a, we pooled each animal’s sessions (within session type). We computed smoothed occupancy maps of the cage (25-pixel bins) and of the behavioral space over the first 30 minutes. The maps show the average difference between stimulus and control sessions across animals. For the mask stimulus, each stimulus session’s cage map was rotated so that the stimulus was at the bottom of the map.

### Test-retest analysis and trait groups

For each animal, the recording periods were divided into two halves, either by alternating between periods or by comparing the earlier periods to the later periods. Baseline features were measured in control sessions (the mean of the control sessions in each period) and stimulus features were measured as the difference between each type of stimulus session and the mean of that period’s control sessions. These analyses used the first 30 minutes of each session. In cases where the stimulus was presented multiple times within a period (such as the toys), we only used data from the first presentation. Breeding pairs and their juvenile offspring were included if they had a full feature set in at least two recording periods. This resulted in 64 animals (35 adults and 29 juveniles) from 19 families. Each feature was z-scored across animals and halves.

We calculated similarity between two animals (or between the two halves of a single animal’s data) using the Pearson correlation across the 10 selected features. This was done using the first half from one animal and the second half from the other animal (or the same animal). Animals were clustered using average-linkage hierarchical clustering of this similarity (distance = 1 - similarity), and the tree was cut into three groups using the alternating halves.

To measure the consistency of the groups, we assigned each half of each animal to a group. For each half, we computed its correlation with the mean profile of each group in the other half. The mean profiles were computed without that animal. The half was assigned to the group with the highest correlation. We then counted the animals whose two halves were assigned to the same group. We compared this count with a null distribution in which the group assignments of the second halves were shuffled across animals. As a further reference, every value of the underlying features was replaced by random noise, and the split, z-scoring, clustering, and assignment were repeated. The association between age and group was tested with a permutation test of the chi-square statistic, shuffling age within families.

### Trait groups within families

For each pair of animals, we determined whether both animals were assigned to the same group. Pairs were classified as cagemates (the two adults in a cage), same-age siblings (pairs of juveniles in the same cage), parent-offspring pairs (an adult and a juvenile in the same cage), or animals of the same age class (two adults or two juveniles) from different families. The number of pairs expected in the same group by chance was calculated from the group sizes within each age class (adults and juveniles), and a 95% chance range (2.5th to 97.5th percentiles of the null distribution) was obtained from 5,000 shuffles of the group labels within each age class.

### Baseline and stimulus dimensions

Baseline features (1,337 features describing behavioral state usage, transitions between states, bout durations, occupancy of the cage and of the behavioral space, movement speed, posture, proximity to cagemates, and time spent in bed during control sessions) were divided into two halves by alternating recording periods. See code for the exact calculation of each feature. From the baseline features, we found that 790 features had values that were correlated across halves (r > 0.5). We considered these features to be “reliable”. The reliable features were grouped into dimensions using the covariance between halves, with the number of dimensions determined by parallel analysis (1,000 permutations). The dimensions were then varimax-rotated, and each feature was assigned to the dimension with its largest loading.

For the scores in Fig. 8c and 8e-f, we repeated this procedure after removing redundant features. To do this, we ranked the reliable features from most to least reliable. We kept each feature in turn unless its true-score correlation with a feature already kept was 0.9 or higher in absolute value. This left 358 features. We then matched each resulting dimension one-to-one to a dimension of the full set, by maximizing the summed absolute true-score correlation between their scores. Each matched dimension was given the same sign as its counterpart in the full set.

Stimulus dimensions were obtained in the same way from the 55 reliable stimulus-response features measured in the tablet, treat, mask, and intruder sessions. See code for exact calculation of each feature. To find the true-score correlation between a baseline dimension and a stimulus dimension, we measured the correlation between one half’s baseline score and the other half’s stimulus score divided by the square root of the product of the two scores’ reliability (their correlations across halves).

To predict each stimulus dimension from baseline behavior, we used nested ridge regression with all 1,337 baseline features. Predictions were made from one half of the baseline data to the other half of the stimulus data, either for families that were held out from training (leave-one-family-out cross-validation) or for animals held out with their cagemates included in the training data (8-fold cross-validation). The ridge penalty was chosen by cross-validation within the training data. The share of reliable variance predicted was reported as the squared correlation between the predicted and observed scores, divided by the reliability of the stimulus score.

### Animal and data selection

The marmoset colony was genetically heterogeneous (unlike mouse colonies that are often inbred) and individual animals differed genetically. They also differed in their housing conditions, such as the room in which their cage was kept and the size and composition of their family unit (i.e. a pair of animals, or two adults with their juvenile offspring). Because data collection in this colony is ongoing, with new controls and experiments added continuously, we analyzed the recordings that were available and sufficiently complete at the time of manuscript preparation. Some analyses applied inclusion requirements, such as a minimum number of control sessions or recording periods per animal. Additionally, a small amount of data was not collected (for example, due to holidays or scheduling conflicts), lost due to human error, or corrupted after being stored for a long period of time. Because of these inclusion criteria and small amount of missing data, the sample size (n) for animals and sessions included differs between figures.

No animal, session, or recording was excluded from any analysis on the basis of its behavioral or vocal measurements or on the basis of any analysis result (i.e. based on how strongly call rate correlated with behavior). Apart from cage and family membership, which defined cagemates and family relationships (Fig. 3e, Fig. 8), the only predefined groups of animals that we compared were based on housing (paired versus family-housed animals; Fig. 4, Fig. S10) and age group (adults versus juveniles; Fig. 7i, Fig. S14). The trait groups (Fig. 7) were not defined in advance but were obtained by clustering the animals’ behavioral data.

### Statistical analysis

Statistical analyses were performed in Python using NumPy and SciPy. Unless stated otherwise, the unit of analysis was the animal, tests were two-sided, and p values were not corrected for multiple comparisons. Tests and confidence intervals across animals treated cagemates as independent, although the behavior of cagemates was correlated (Fig. 3e). We examined the nature and extent of those correlations and described them (Fig. 3 and Fig. 4).

### Comparisons between conditions and age groups

We compared control and tablet sessions with paired t-tests (Fig. 5; n = 72 animals, 70 for discriminability). We also compared stimulus and control sessions with paired t-tests (Fig. S13; n = 68-92 animals per condition). Each test used one mean value per animal in each condition. In Fig. S14, we compared juvenile and adult sessions with Student’s t-test. This test included 1,877 sessions from 49 juveniles and 2,486 sessions from 47 adults. We compared juveniles and adults that were recorded together with a paired t-test (1,690 pairs). Because sessions from the same animal are not independent, we also compared the family medians of the juvenile and adult values (paired t-test, n = 31 families).

### Correlations and discriminability

In Fig. 3e, we correlated each animal’s values with the mean of its cagemates’ values within a recording period (n = 331 animal-periods from 96 animals). We compared these correlations with a reference. For the reference, we correlated the same animal’s values with the cagemate mean from one randomly drawn cage-period. This cage-period came from another cage or from the same cage in another recording period. The two sets of values were aligned by session type. Both measures are shown as means with t-based 95% confidence intervals. The correlations between call rate and behavior (Fig. 4, Fig. S10) were Spearman correlations across 5-minute segments within each animal. The two animals of a pair shared one call-rate series (68 animals, 35 pairs). Family-housed cagemates that were recorded together also shared a call-rate series.

Each AUROC (Fig. 6c, e; Fig. S12) was the fraction of all combinations of animals in which the projection in the first condition was larger than the projection in the second.

### Trait groups

We compared the agreement between halves (Fig. 7b-c) with a null distribution. In each permutation, the group assignments of the second halves were shuffled across animals. This kept the size of each group constant. We used 2,000 permutations for alternating halves and 20,000 permutations for early/late halves. For adults and juveniles, the assignments were shuffled within each age class.

We also measured the agreement that the procedure produced from random data. In each of 200 draws, every value of the underlying features was replaced by noise from a standard normal distribution. We then repeated the split, z-scoring, clustering and assignment with the same 10 features.

### Families and dimensions

For each pair type (Fig. 8b), we computed the number of pairs expected to share a group by chance. This expectation was computed exactly from the group sizes within each age class. The 95% chance range (2.5th to 97.5th percentiles) came from 5,000 permutations. In each permutation, the group labels were shuffled among all adults, and separately among all juveniles.

We used parallel analysis to choose the number of baseline and stimulus dimensions. We generated 1,000 permuted data sets. In each one, animals were shuffled independently for each feature. The same shuffle was used in both halves. We kept components in order while each eigenvalue was larger than the 95th percentile of the eigenvalue of the same rank in the permuted data. In Fig. 8e-g, each animal contributed two points. One point compared its first-half score with its second-half score, and the other point compared them in the reverse direction. The correlation was therefore computed over 128 points.

We tested the share of reliable variance predicted from baseline features with 200 permutations. We did this for the held-out families (Fig. 8d) and for the 8-fold cross-validation. In each permutation, we shuffled the stimulus scores either across all animals or within each family and age class (for example, among the juveniles of one family). We then reran the whole nested procedure, including the choice of the ridge penalty. Each permutation used 4 repeats of the 8-fold partition. The observed data used 20 repeats. For Fig. 8, p was computed as (1 + k)/(1 + N) where k is the number of permutations that reached the observed value, and N is the total number of permutations.

### Data and code availability

The source data and code needed to reproduce each figure panel are available at: https://github.com/williammenegas/Marmoset_Quantification_2026. Processed data types such as postural time series data, behavioral state data, and vocalization time series data are publicly hosted on Zenodo. The marmoset behavioral analysis described here is produced by several related code repositories. marm_behavior (https://github.com/williammenegas/marm_behavior) is the behavioral pipeline running pose estimation, per-animal body-part tracking, per-frame feature extraction, and projection of those features into a behavioral cluster space. marm_filt (https://github.com/williammenegas/marm_filt) denoises the body-part tracks before use, applying a graph neural network with temporal convolutions implemented in TensorFlow and trained on the tracks. marm_viz (https://github.com/williammenegas/marm_viz) is a general-purpose viewer for the output files, providing the spatial occupancy, density, ethogram, state-usage, t-SNE coordinates, head-direction, and transition-matrix views.

## ACKNOWLEDGEMENTS

We thank Alex Mathis and Mackenzie Mathis for the helpful discussions and access to beta versions of their software. We thank the MIT Division of Comparative Medicine (DCM) for support with animal care.

## ETHICS STATEMENT

All marmoset procedures were performed in accordance with the National Institutes of Health Guide for the Care and Use of Laboratory Animals and were reviewed and approved by the Massachusetts Institute of Technology Committee on Animal Care (CAC), MIT’s Institutional Animal Care and Use Committee, under protocol 2303000479. This protocol (“Genetic manipulation of common marmosets for the advancement of neuroscience research”, with Principal Investigator: Guoping Feng) has been continuously approved across renewal cycles for the duration of this study. MIT holds Public Health Service Animal Welfare Assurance D16-00078 and USDA registration 14-R-0018. Animals were housed, monitored, and provided veterinary care by the MIT Division of Comparative Medicine.

## AUTHOR CONTRIBUTIONS

W.M. and E.C. designed the study. W.M., E.C., S.P., H.X., and K.B. collected the behavioral data. W.M. performed analysis and wrote the manuscript. G.F. and R.D. supervised the project. All authors reviewed the manuscript.

---

**Figure S1:**
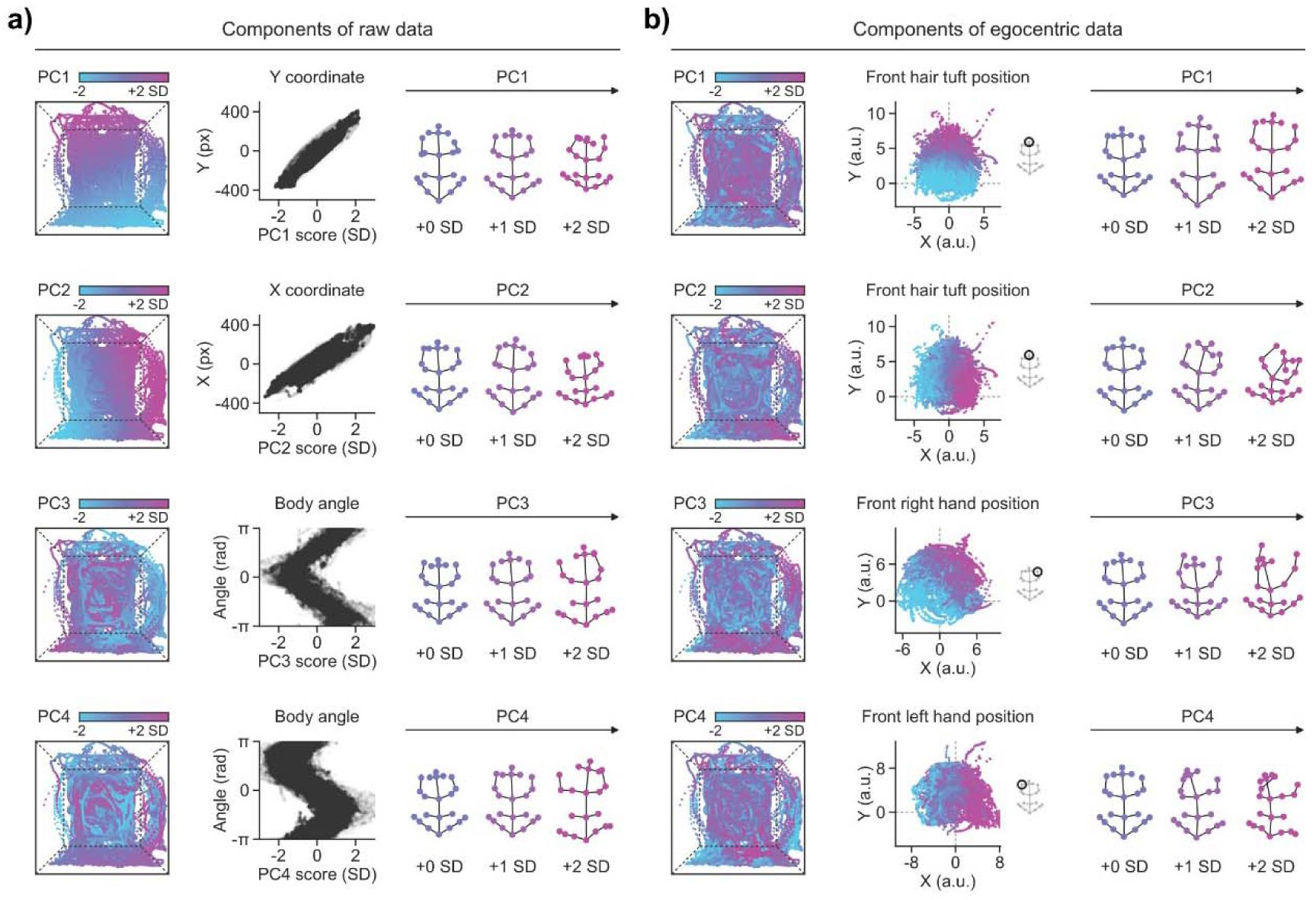
PCA in raw data compared to egocentric data. **a)** Illustration of the first 4 principal components (PCs) of the raw postural data (the X and Y coordinates of 21 keypoints). Left: head position in the cage colored by PC score. Middle: PC score plotted against X coordinate, Y coordinate, or body angle. Right: Average egocentric posture at +0, +1, and +2 standard deviations (SD) of the score. PCA was computed over 60 minutes of tracking data. **b)** Illustration of the first 4 principal components (PCs) of the egocentric postural data (normalized by position, body angle, and body length). Left: head position in the cage colored by PC score. Middle: the egocentric position of a keypoint colored by score. Right: average egocentric posture at +0, +1, and +2 SD of the score. PCA was computed over 60 minutes of tracking data. Dashed lines show a schematic of the cage.

**Figure S2:**
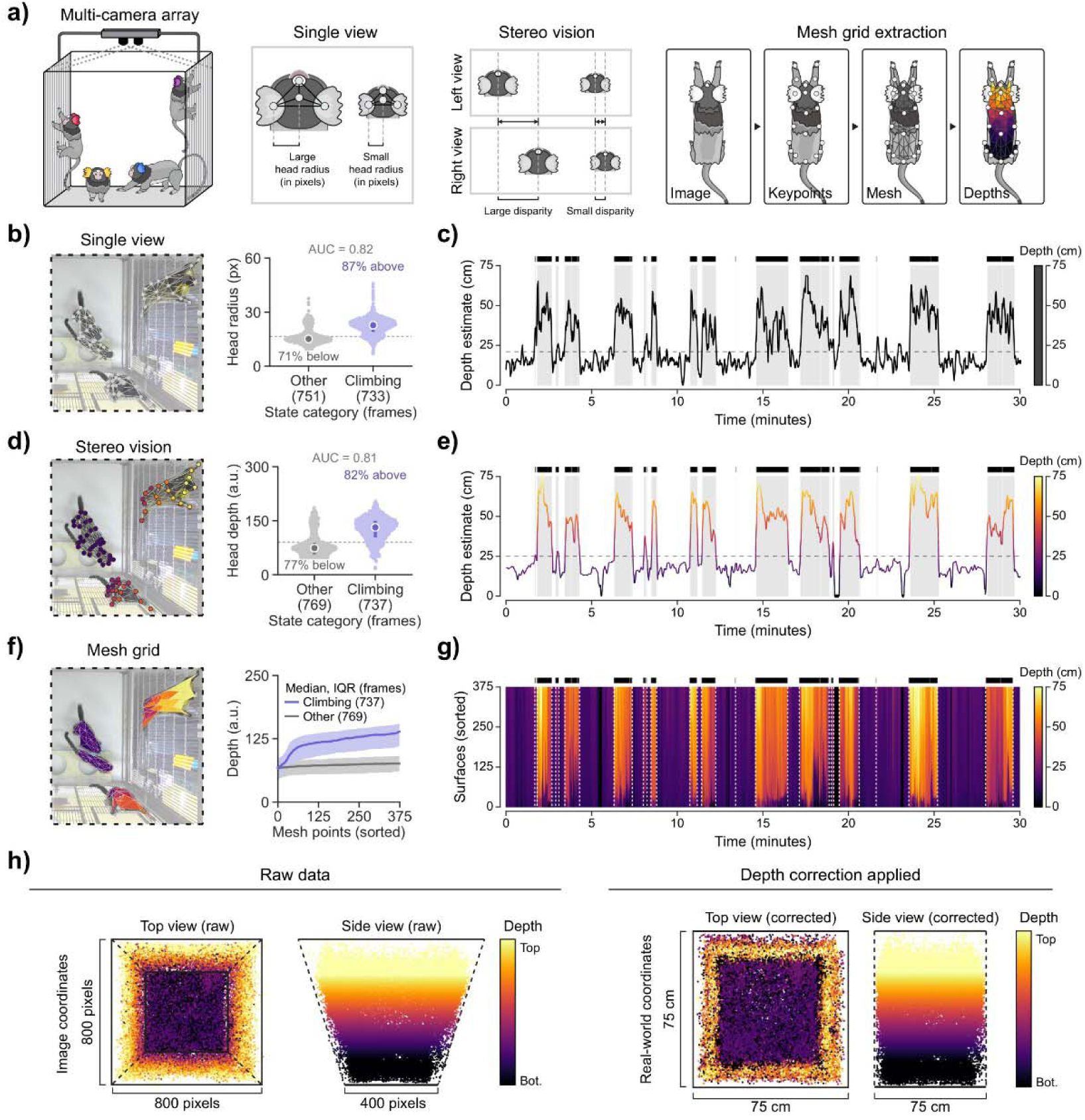
Depth estimates for top-down home-cage video. **a)** Comparison of methods for depth estimation using a single view, stereo vision, or stereo vision with mesh grid extraction. **b)** Example single-view image, and head radius (pixels) in frames whose behavioral state is in the climbing category (n = 733 frames) and in all other states (n = 751 frames). The dashed line indicates the threshold that best separates the two groups. **c)** Example depth time series based on single view (30 minutes). Shading and black bars indicate climbing-category bouts, and the dashed line indicates the threshold from single-view depth estimation. **d)** Example stereo vision image, and head depth from the stereo depth map (a.u.) for climbing-category frames (n = 737) and all other frames (n = 769). **e)** Example depth time series based on stereo vision from the same 30-minute window, colored by depth. **f)** Example mesh grid based on stereo depth estimates, and depths at 375 mesh points around the animal’s outline. **g)** Example mesh grid time series from the same 30-minute window, with depths sorted at each time point. **h)** Head positions from one session before (left) and after (right) depth correction, viewed from the top and from the side. Color indicates estimated head depth.

**Figure S3:**
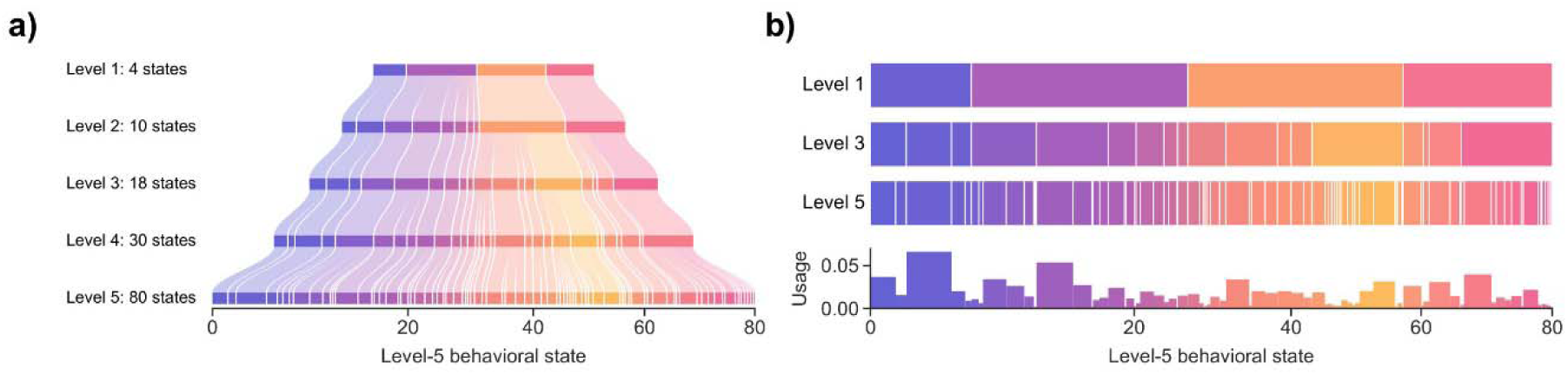
Hierarchical behavioral state representation. **a)** Usage of behavioral states across the five hierarchical levels (comprised of 4 clusters, 10 clusters, 18 clusters, 30 clusters, and 80 clusters). Data was pooled over a random sample of 250 sessions. The size of each block indicates the fraction of time spent in that behavioral state, and ribbons show the correspondence between clusters at different hierarchical levels. **b)** The same hierarchy shown at levels 1, 3, and 5. Below: bar chart of the usage of each state. Colors indicate behavioral category.

**Figure S4:**
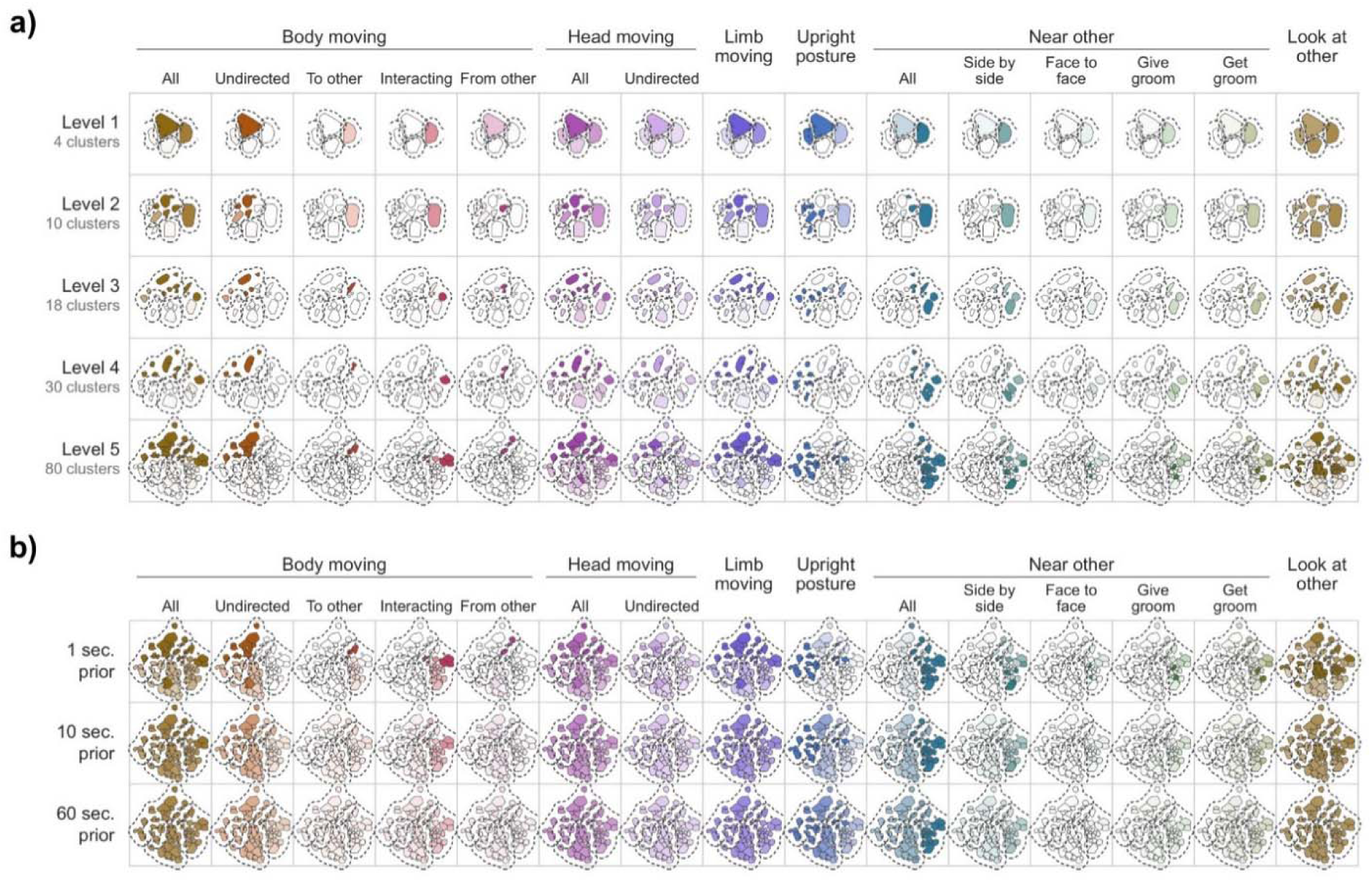
Mapping human description and features to clusters in latent space. **a)** Distribution of selected behavioral variables (binary per-frame features computed from pose and depth) across the behavioral latent space at each hierarchical level. Each column corresponds to a behavioral feature, and each row corresponds to a hierarchical level. The cluster outlines are shaded based on the share of each cluster’s frames that display the feature. Dashed lines outline the behavioral categories. **b)** Distribution of behavioral variables at the lowest hierarchical level for the frames 1 second (top row), 10 seconds (middle row), and 60 seconds (bottom row) before the labeled frame. These distributions show persistent structure in the data.

**Figure S5:**
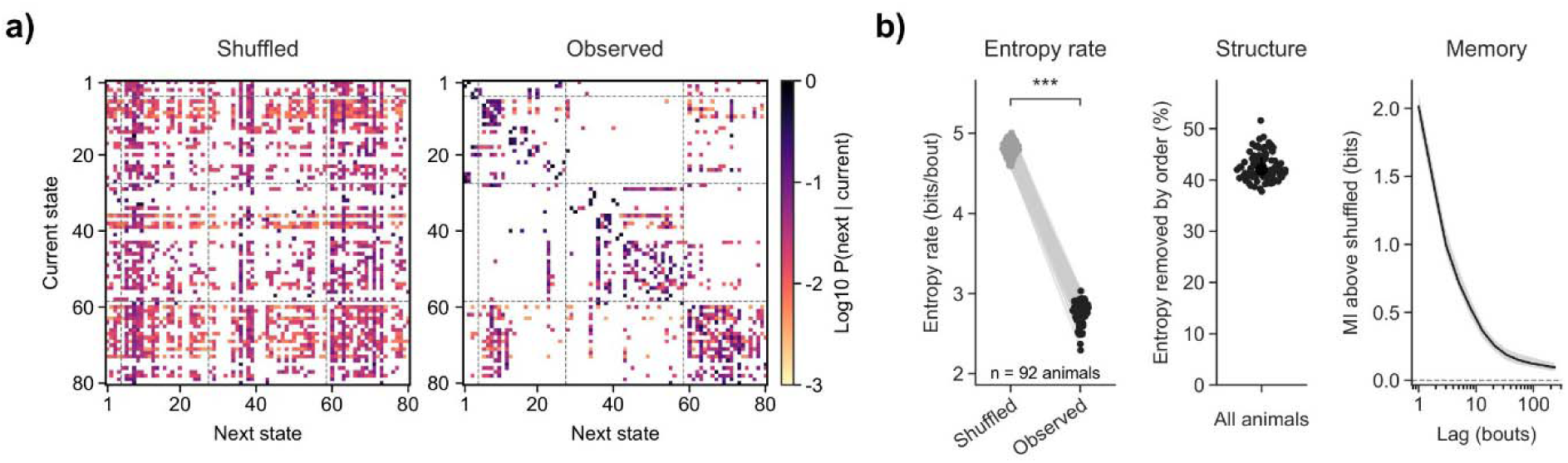
Transition structure of behavioral state sequences. **a)** Transition probabilities between behavioral states for an example session (comprised of 4864 behavioral state transitions), comparing the shuffled sequence (left) and actual sequence (right). Dashed lines indicate boundaries between behavioral categories. **b)** Left: entropy rate of behavioral state sequences (bits per bout) for shuffled and observed sequences. Middle: the percentage of entropy removed by the order of states. Right: the mutual information between states that are separated by an increasing number of bouts. The median, quartiles, and individual data are plotted. *** p < 0.001.

**Figure S6:**
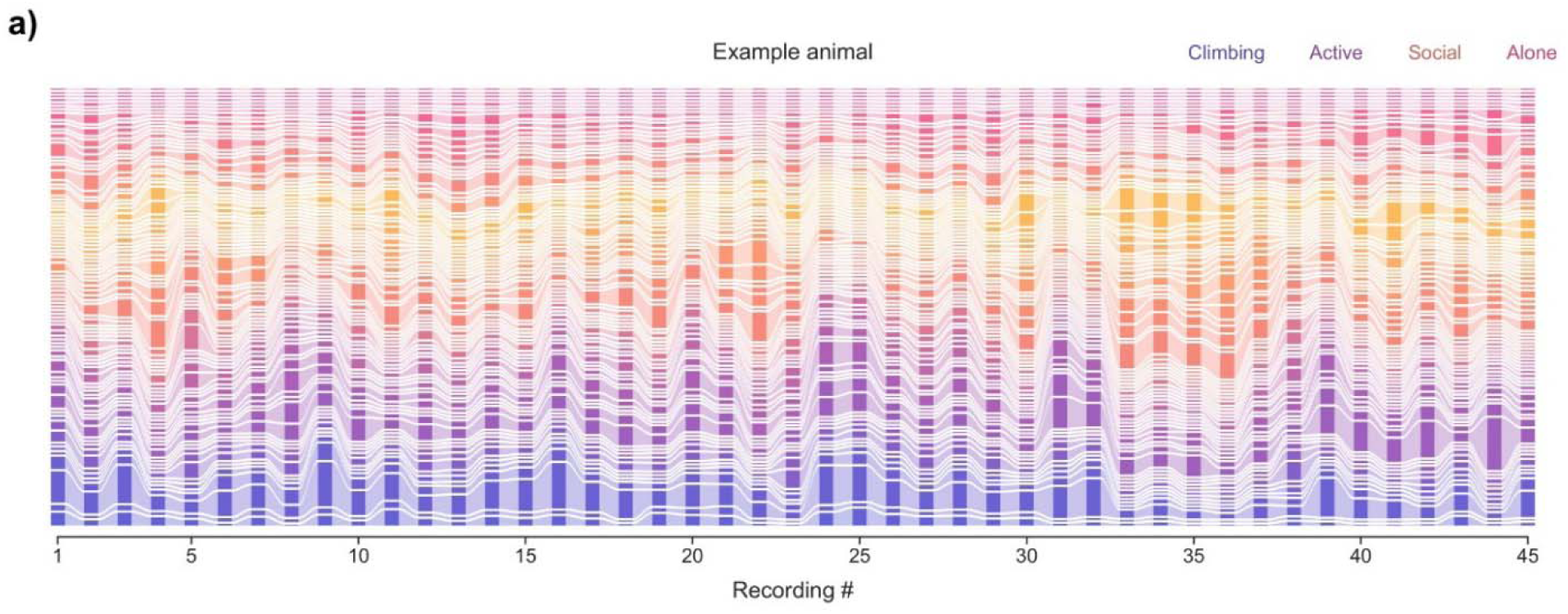
Behavioral state usage across sessions of one animal. **a)** The usage of 80 behavioral states across 45 recordings from one example animal. Data from multiple non-consecutive recording periods are concatenated. Band height indicates the fraction of labeled frames in each state, and band colors indicate behavioral category.

**Figure S7:**
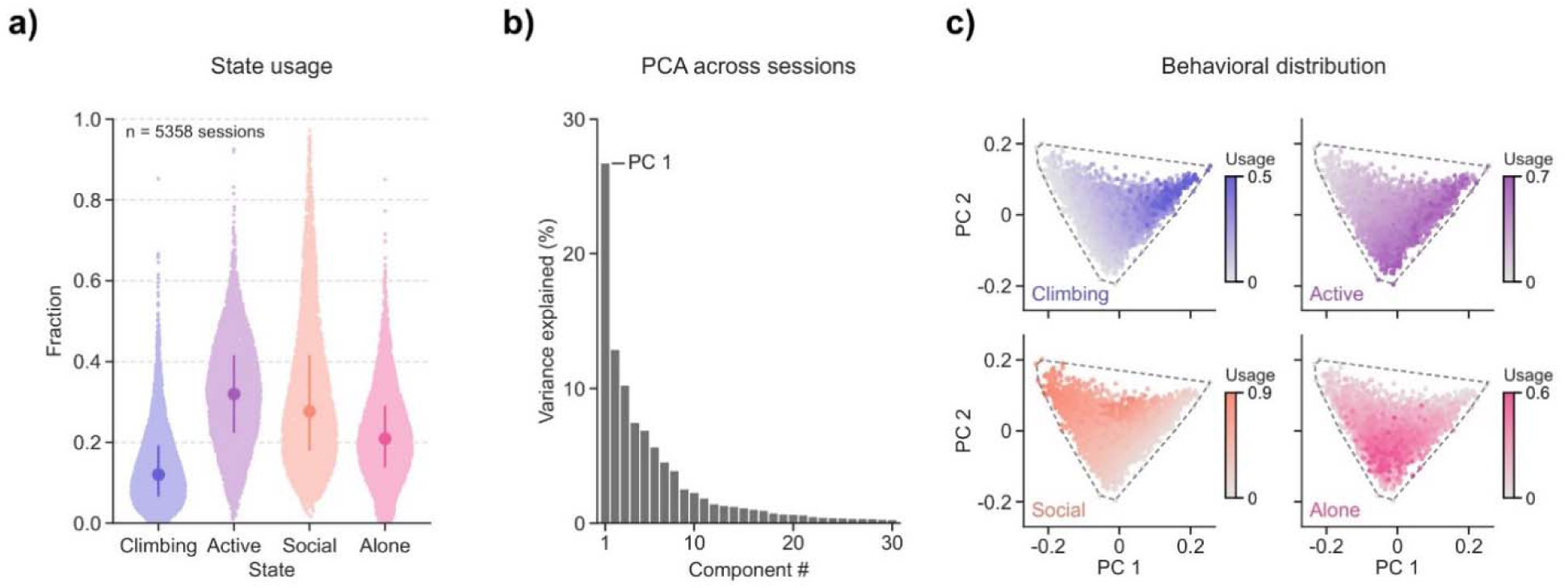
Behavioral state usage and PCA across sessions. **a)** Fraction of each session spent in 4 coarse behavioral categories across 5358 sessions from 116 animals, with each point representing one session. The median, quartiles, and individual data are plotted. **b)** Variance explained by each component of a principal component analysis across sessions based on the usage of the 80 fine behavioral states. **c)** All sessions plotted by PC 1 and PC 2. Scattered points are colored based on the fraction of time spent in each behavioral category.

**Figure S8:**
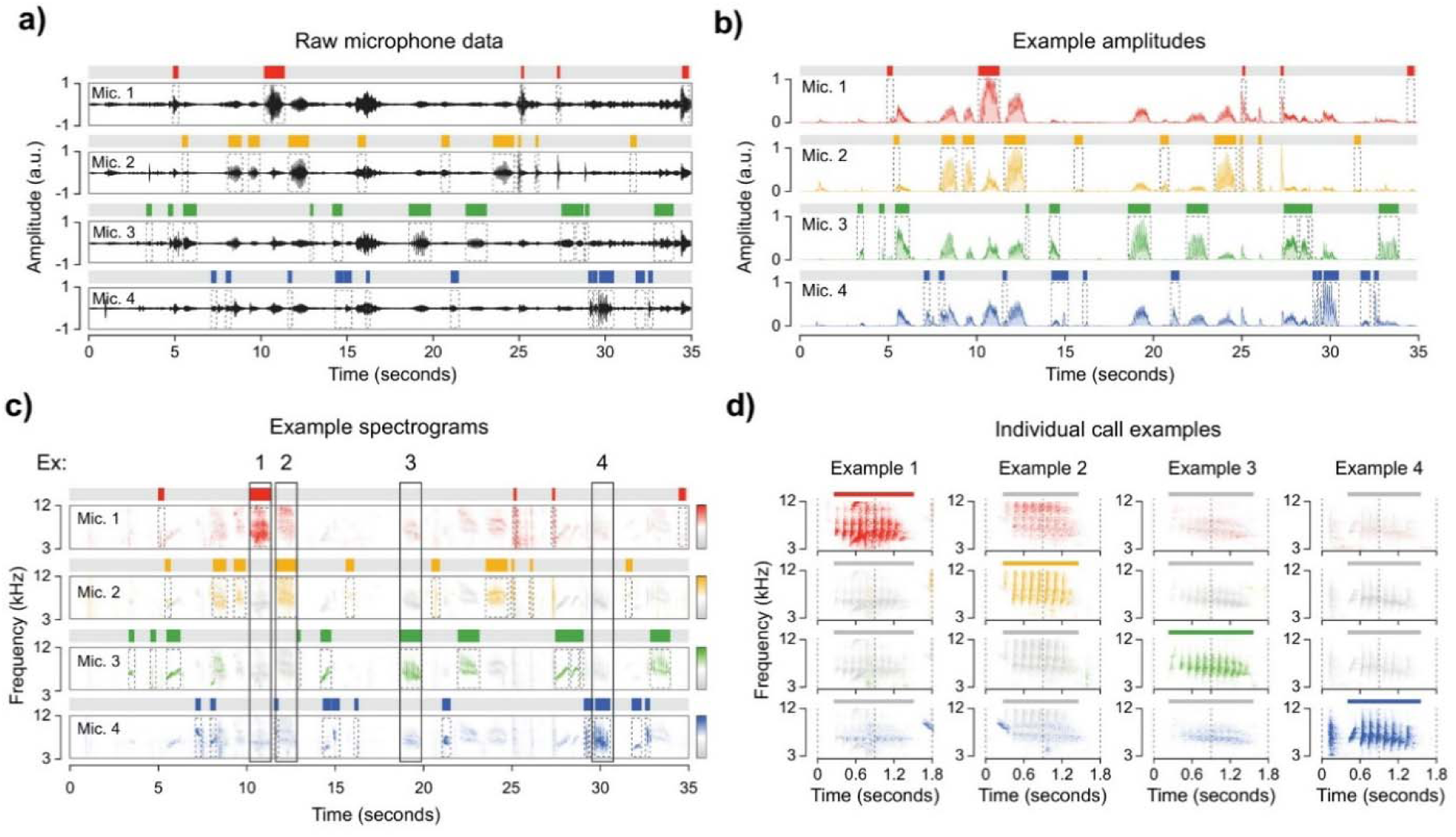
Call localization using an array of microphones. **a)** Example time series (35 seconds) of raw data from 4 microphones placed near different cages. Dotted boxes and colored bars indicate call timings labeled by a neural network. The call is attributed to a microphone based on amplitude. **b)** Example time series of amplitudes (running average of the absolute value of the spectrogram band) used to assign each call to the microphone with the largest amplitude. **c)** Example spectrograms (3-12 kHz) of the same 35 seconds as in other panels, showing the distribution of signal across frequencies. **d)** The spectrogram from four individual calls, with brightness indicating the amplitude of the call on each microphone relative to the other microphones.

**Figure S9:**
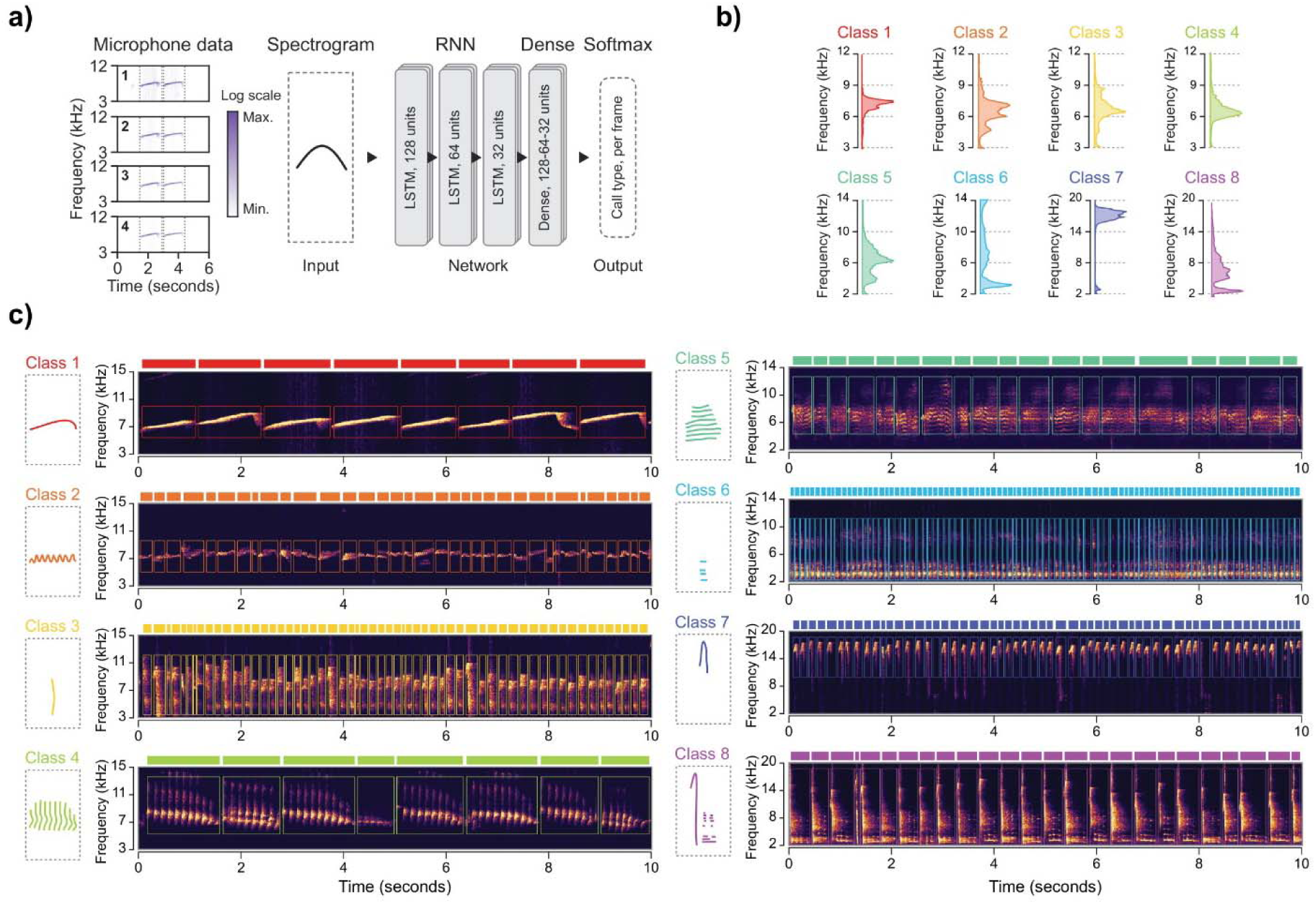
Supervised categorical audio classification. **a)** Schematic depicting the strategy for classifying calls by call type using a supervised neural network. Spectrograms from each microphone are passed to a recurrent network that outputs a call type for each frame. **b)** Frequency distribution of each call type, colored by call type. **c)** Left: human-drawn representation of each call type. Right: example calls (10 seconds of concatenated non-consecutive calls) from each type, chosen randomly from all calls of that type. Boxes and bars show the onset and boundaries of individual calls. The number of calls displayed here varies by type because call duration differs across types. Classes were defined by human categorization of calls in the training set and not named.

**Figure S10:**
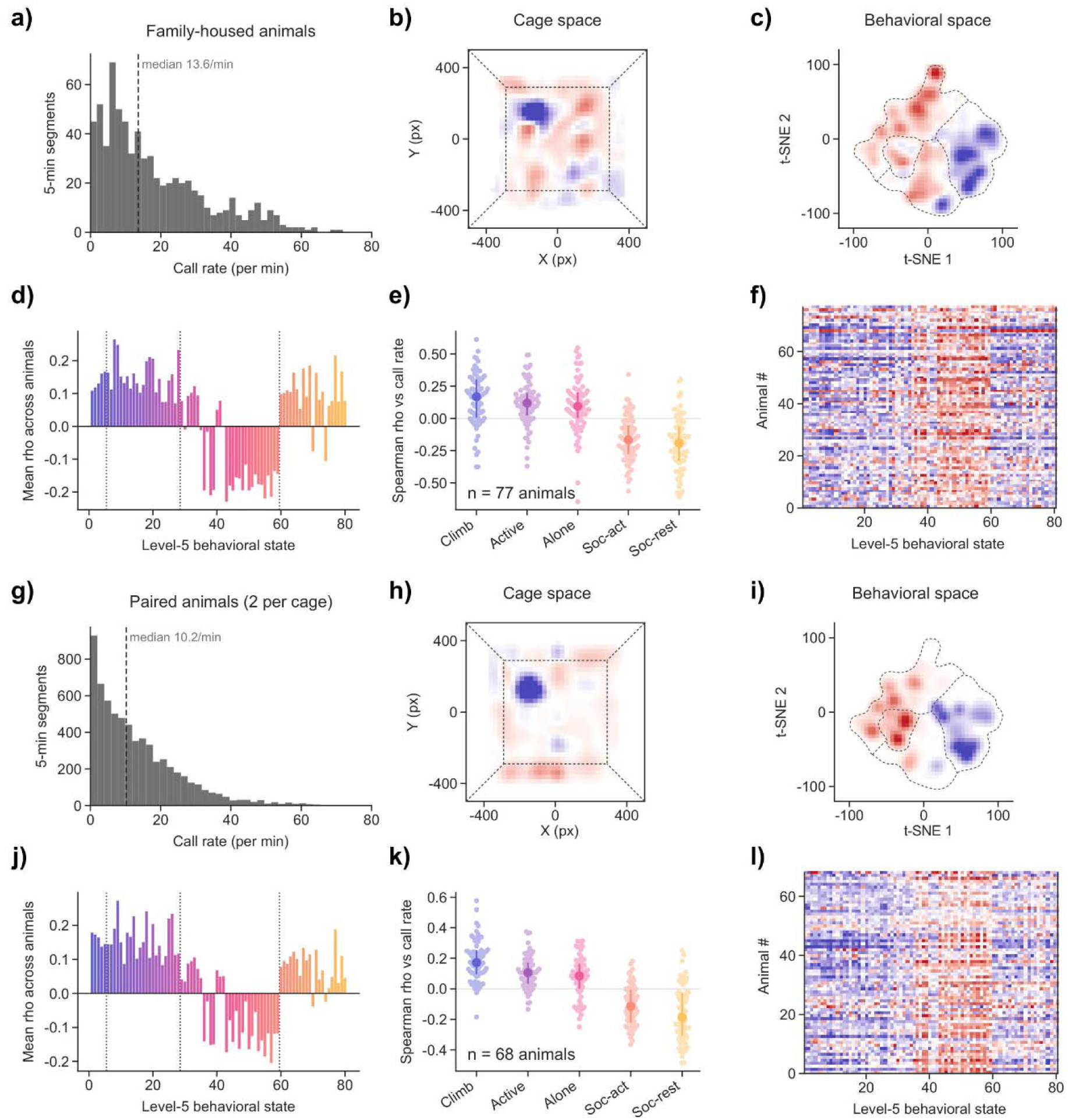
Comparison of vocalization-related behavioral patterns across family sizes. **a)** Distribution of call rates in 5-minute segments from control sessions of family-housed animals (n = 702 segments), with the dashed line indicating the median. **b)** Difference in cage occupancy between segments with call rates above and below each recording’s median, averaged across animals (n = 81 animals). **c)** The same comparison in behavioral embedding space. **d)** Average Spearman correlation between call rate and the usage of each behavioral state across animals. **e)** Correlation between call rate and behavioral category usage. The median, quartiles, and individual data are plotted. **f)** Spearman correlation between call rate and the usage of each behavioral state for each animal. **g-l)** The same analyses for pair-housed animals (two per cage) from n = 68 animals.

**Figure S11:**
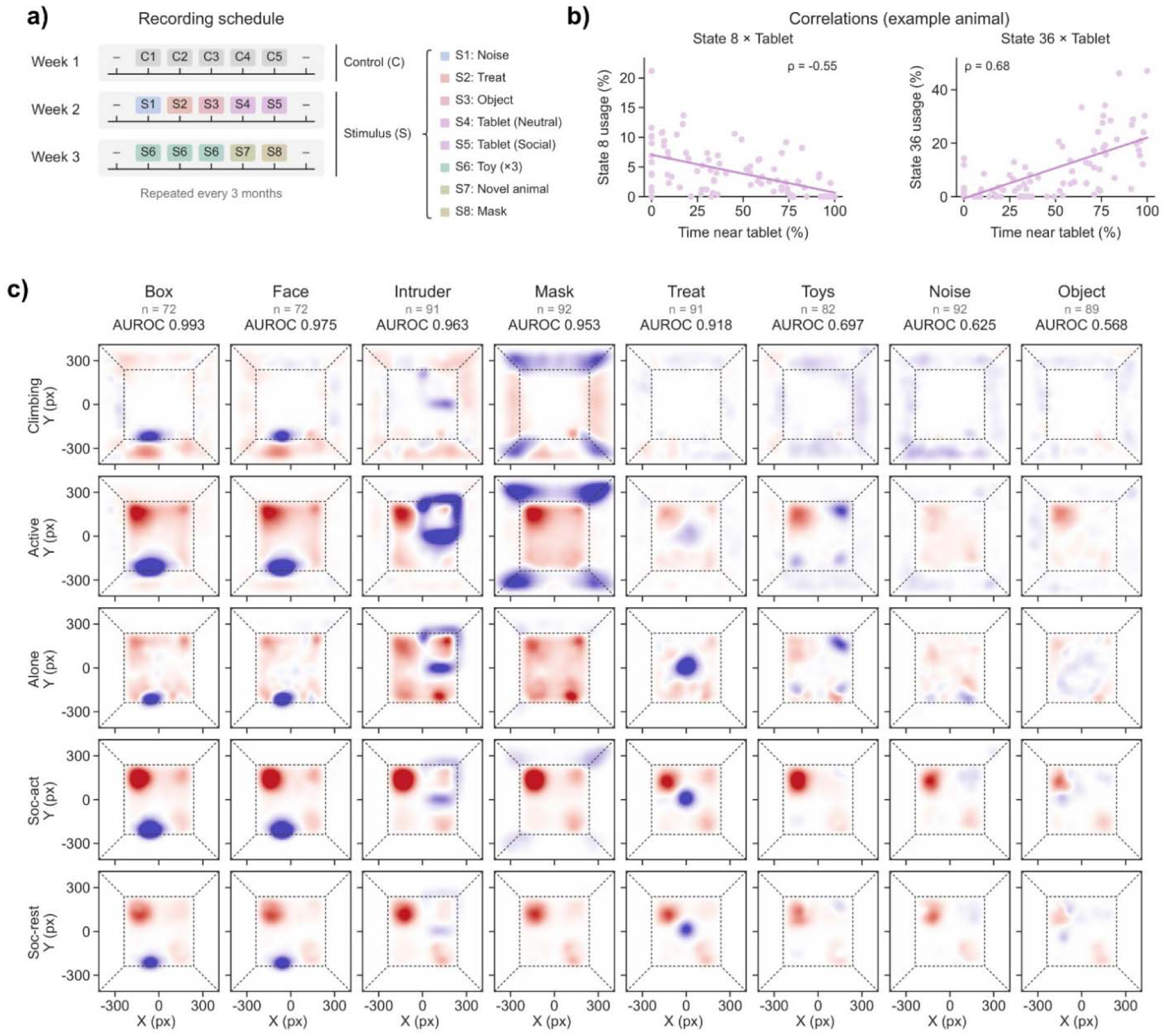
Behavioral responses to a panel of stimuli. **a)** Recording schedule. The first week contains five control sessions (C1-C5). The second week contains generally appetitive stimuli (treat, novel object, and a tablet delivering syrup reward). The third week contains toys, and also highly salient stimuli such as the novel animal and the mask. **b)** Example correlations between behavioral state usage and time spent near the tablet in one animal, using 1-minute bins. ρ indicates Spearman correlation. **c)** Average change in cage occupancy elicited by each stimulus relative to control sessions. Each row shows the change in occupancy specific to one behavioral category. Data is pooled across the first 30 minutes of each session. Blue indicates locations used more during the stimulus session and red locations used less. The number of animals (n) and the discriminability of each animal’s response from its held-out control session (AUROC) are listed above each column.

**Figure S12:**
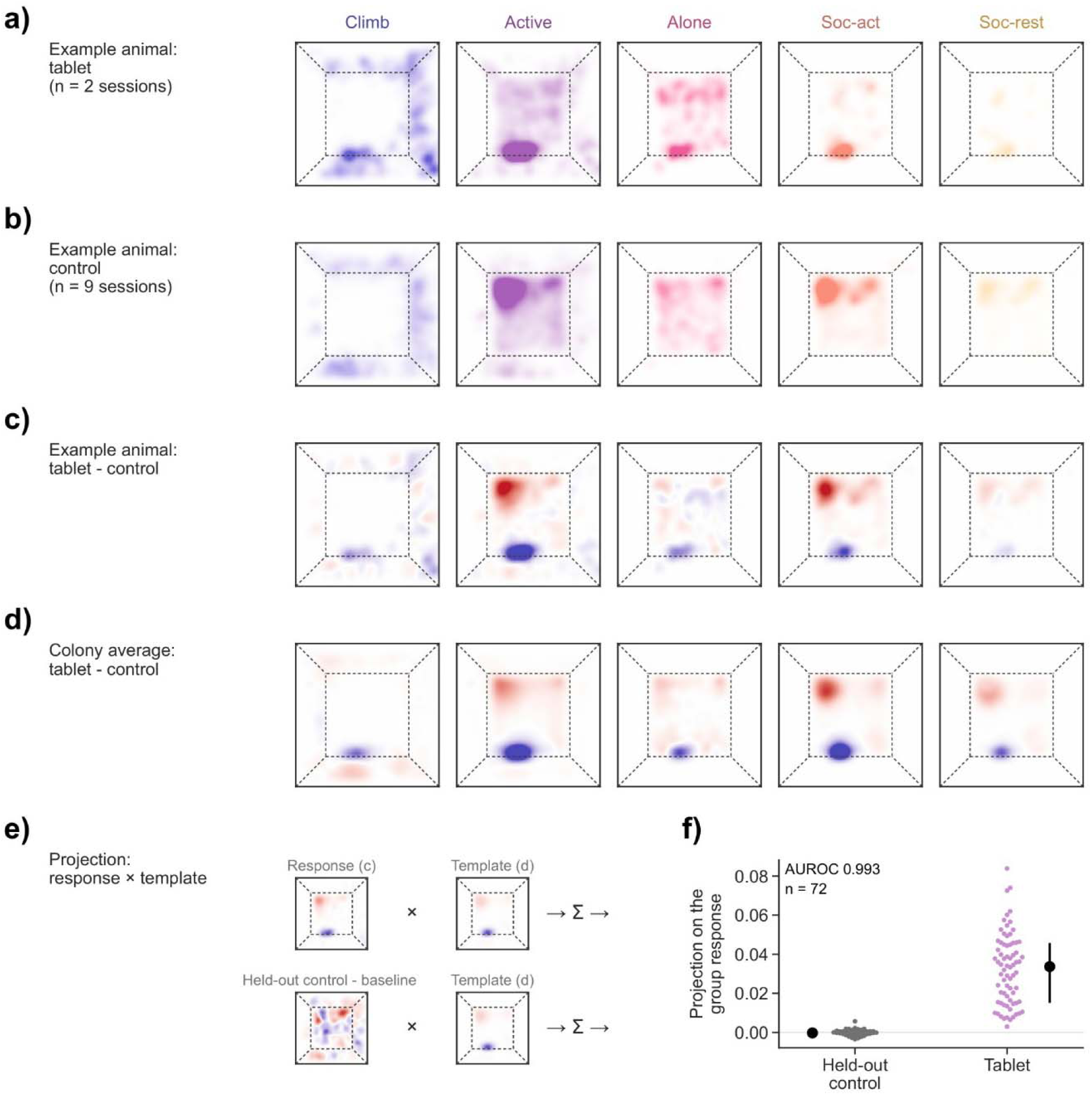
Measuring stimulus responses by projection on the group response. **a)** Cage occupancy maps in each behavioral category over the first 30 minutes of the tablet sessions of an example animal. **b)** The same maps averaged across control sessions from that same example animal. **c)** The difference between each set of maps. Blue indicates higher occupancy when responding to the tablet and red indicates lower occupancy. **d)** Average response across the colony used to assess similarity. **e)** Schematic of the projection (see Methods for details). **f)** Projection on the group response for the held-out control session and the tablet session for each animal (n = 72 animals). The AUROC is the mean across 10 draws of the held-out control. The median, quartiles, and individual data are plotted.

**Figure S13:**
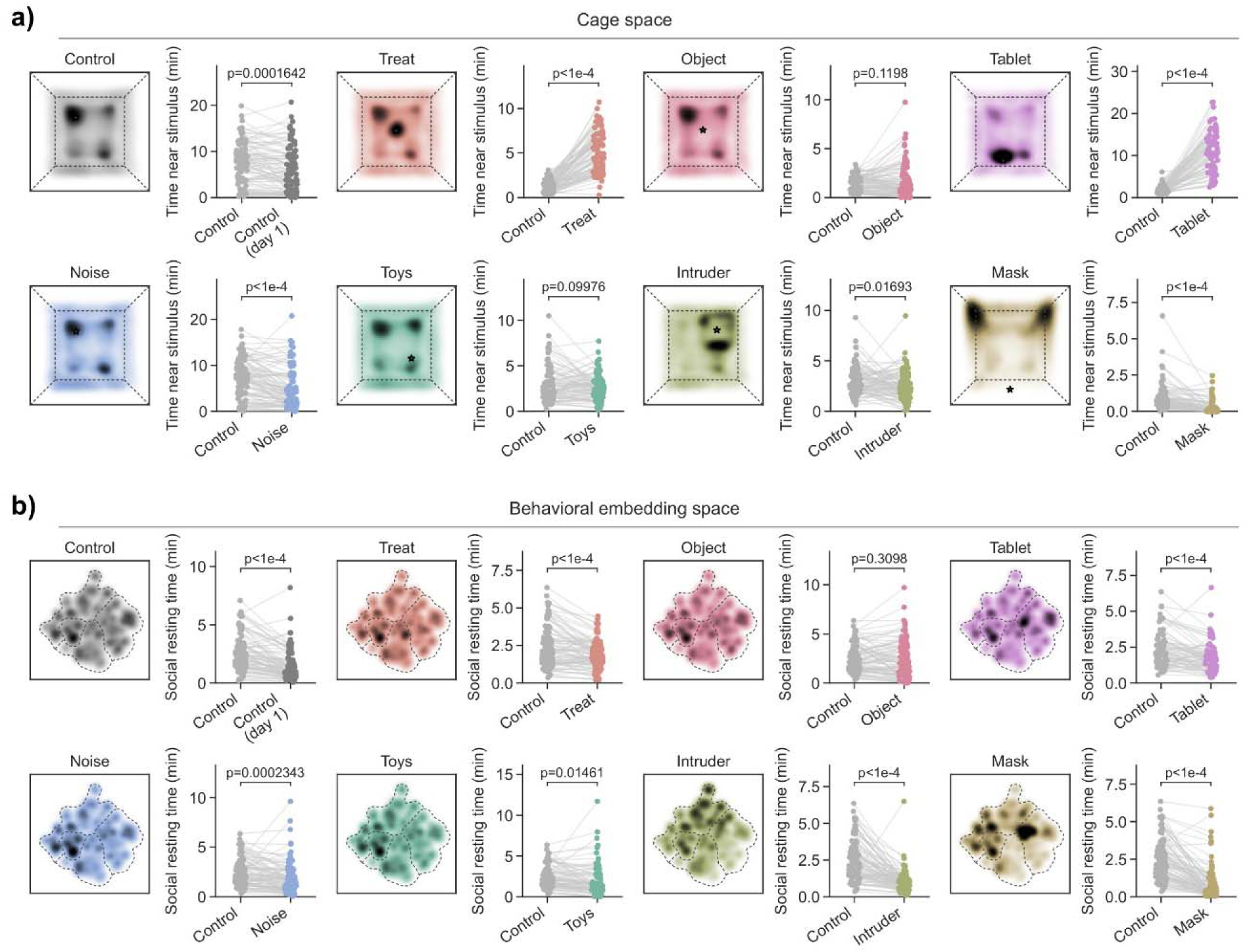
Time near stimulus and social resting time for each stimulus condition. **a)** Average cage occupancy over the first 30 minutes for each condition, with marker indicating the stimulus location. Right: time spent near the stimulus (over the first 30 minutes) in control and stimulus sessions for each animal. For the control condition, the first control session at each age (control day 1) is compared with the animal’s other control sessions. The mask map is rotated so that the stimulus is at the bottom. The p-values shown are from two-sided paired t-tests. **b)** Average occupancy of behavioral embedding space for each condition (dotted lines outline the behavioral categories). Right: time spent in social resting states in control and stimulus sessions for each animal. The p-values shown are from two-sided paired t-tests.

**Figure S14:**
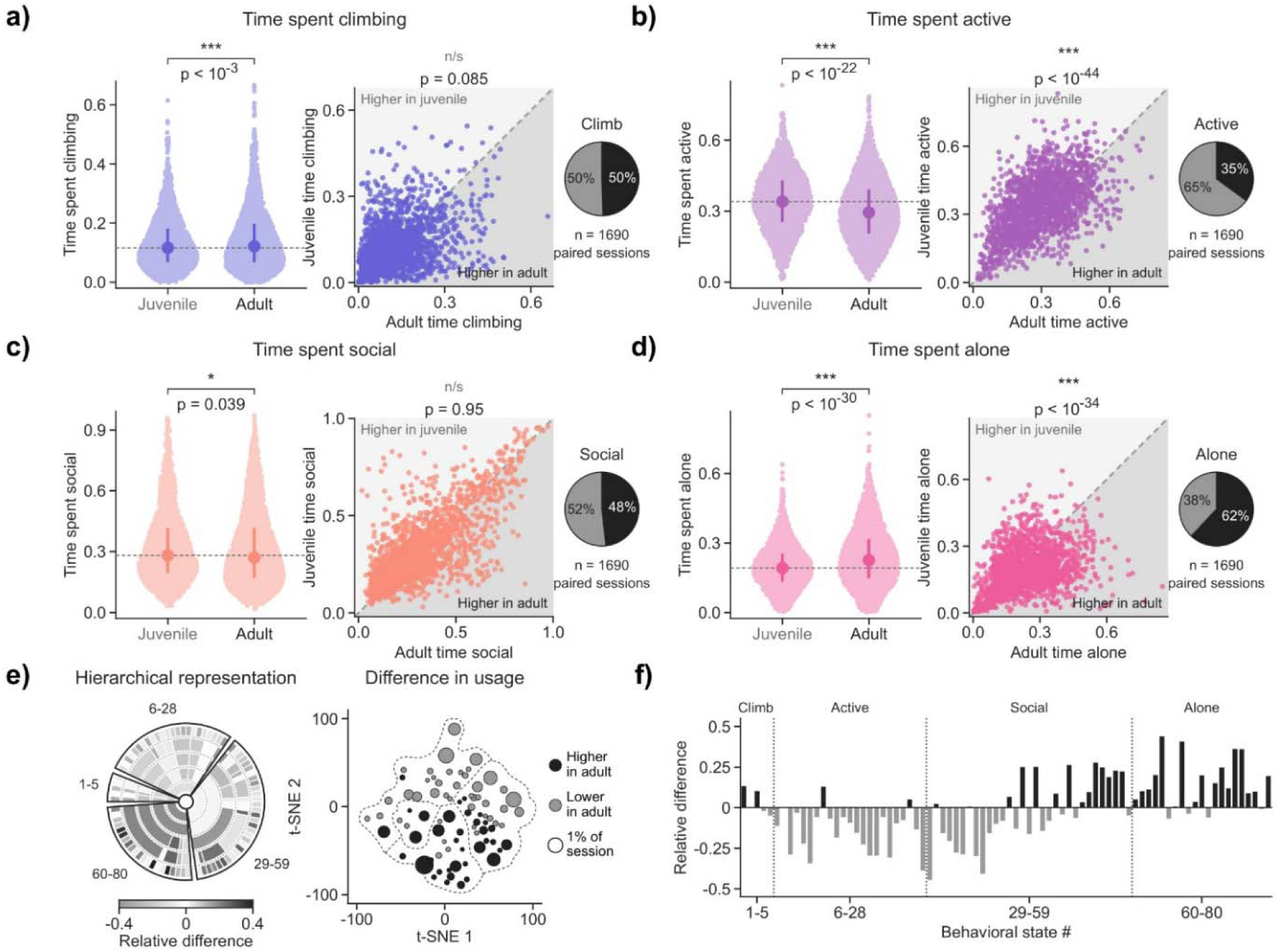
Differences in behavioral distribution between juvenile and adult animals. **a)** Fraction of time spent climbing per session in juvenile and adult animals (left; unpaired t-test), and a paired comparison between cagemates of different ages recorded in the same session (right; paired t-test). Pie chart indicates the fraction of pairs with a higher value in the juvenile (grey) or in the adult (black). **b)** The same comparisons for time spent active. **c)** The same comparisons for time spent social. **d)** The same comparisons for time spent alone. **e)** Relative difference in the usage of behavioral states between adult and juvenile animals. Left: every node of the five hierarchical levels. Right: each behavioral state at its average position in embedding space (black: higher in the adult, grey: lower in the adult). The area of each circle indicates the absolute difference in usage between juveniles and adults. Dashed lines outline the behavioral categories. **f)** Relative difference in usage of each fine behavioral state. Dotted lines indicate boundaries between behavioral categories. * p < 0.05, ** p < 0.01, *** p < 0.001.

